# *In vivo* generation of post-infarct mouse cardiac muscle by cardiomyocyte progenitors produced with a reproducible laminin-promoted human stem cell differentiation system

**DOI:** 10.1101/478883

**Authors:** Lynn Yap, Jiong-Wei Wang, Aida Moreno-Moral, Li Yen Chong, Yi Sun, Nathan Harmston, Xiaoyuan Wang, Suet Yen Chong, Miina K. Öhman, Heming Wei, Ralph Bunte, Sujoy Gosh, Stuart Cook, Outi Hovatta, Dominique P.V. de Kleijn, Enrico Petretto, Karl Tryggvason

## Abstract

Regeneration of human heart muscle after injury is extremely limited and an unmet clinical need. Despite extensive research, there are no methods for the reproducible generation of clinical quality stem-cell-derived cardiovascular progenitors (CVPs). Here, we identified laminin-221 (LN-221) as the main cardiac laminin, which was produced here as human recombinant protein, and showed that LN-221 promotes differentiation of pluripotent hESCs towards cardiomyocyte lineage and downregulates genes associated with pluripotency and teratoma development. We developed a chemically defined, xeno-free laminin-based cardiomyocyte differentiation protocol to reproducibly generate CVPs that form human muscle *in vivo*. We assessed the reproducibility of the differentiation protocol using time-course bulk RNA sequencing developed from two different hESC lines. Single-cell RNA sequencing of CVPs derived from two hESC lines further showed high reproducibility and identified three main progenitor subpopulations. These CVPs were transplanted into myocardial infarction mice, where heart function was measured by echocardiogram and human heart muscle bundle formation was identified histologically. This method may provide clinical quality cells for use in regenerative cardiology.

## INTRODUCTION

It is established that regeneration of human heart muscle after injury is extremely limited ^1,2^. This has led to extensive research efforts aimed at the development of cell therapies for myocardial damage. In particular, multiple studies with adult bone marrow stem cells injected into the infarct region have been carried out, but that approach has failed ^3,4^. Regenerative cardiology needs novel alternative methods for generation of human clinical quality cells that can form new functional cardiac muscle *in vivo*. Numerous reports have described the treatment of heart disease in animals using cardiomyocytes (CMs) derived from human induced pluripotent stem cells (iPSC) ^5^, embryonic stem cells (ESC) ^6–10^ and CMs reprogrammed directly from non-myocytes (induced cardiomyocytes) ^11,12^.

However, despite remarkable recent advances, reproducible methods for the generation of clinical quality cells are still a major challenge. According to regulatory agencies, such as FDA and EMA ^13,14^, therapeutic stem-cell-derived cells are considered as drugs that should be prepared using chemically defined and xeno-free cell culture systems. The cells should also display phenotypic purity and yield reproducible tissue regeneration efficacy and absence of adverse side effects. To that end, a major problem has been that transplantation of human pluripotent stem cell (hPSC)-derived beating CMs leads to arrhythmia and thereby compromises heart function ^8,10^. This questions the applicability of such cells to regenerative cardiology ^15,16^, and suggests that multipotent cardiovascular progenitors (CVPs) might be a possible solution to that problem if they mature to beating CMs *in vivo*.

Another significant problem has been that the differentiation protocols hitherto developed have neither been fully human nor chemically defined, and their reproducibility has varied extensively. For example, the use of an undefined mouse tumor extract (Matrigel™)^17,18^, usually used together with an apoptosis inhibitor (Rho kinase inhibitor), and different animal or human sera render the differentiated cells highly variable and inappropriate for clinical application ^6^. As the ultimate goal of stem cell based therapies is to replace damaged tissue efficiently and safely, it is essential to develop reproducible, defined and xeno-free differentiation protocols for making clinical quality cells that can generate new functional cardiac muscle *in vivo*.

There is evidence that extracellular matrix (ECM) components play a role in cellular differentiation *in vivo* ^19^. However, there is limited knowledge about which specific matrix proteins actually contribute to cell differentiation. Attempts to replace Matrigel^TM^, RGD-containing peptides, recombinant E-cadherin or Synthemax^®^ have not been investigated in *in vivo* studies therefore their success is yet determined ^20–22^.

Basement membranes (BMs) are ultrathin extracellular structures that are located in the immediate vicinity of the plasma membrane of all organized cells ^23,24^. BMs contain structural proteins such as type IV and XVIII collagens, the proteoglycans perlecan and agrin, as well as laminins that exist in multiple, largely cell-type-specific, isoforms. These proteins are bound to the plasma membrane via specific receptors such as integrins. We have produced and extensively studied different BM laminin isoforms, their effects on various cell types and stem cell differentiation *in vitro* ^23,25–29^. Laminins are large trimeric BM proteins that exist in at least 16 isoforms in mammals. Each trimer contains one *α*-, one *β*- and one *γ*-chain that exist in five, four and three genetically distinct forms, respectively. The trimers are named according to their chain composition, *e.g*. the composition of LN-521 is *α*5:*β*2:*γ*1. Laminins are important for cell adhesion and they influence cell differentiation, migration and phenotype stability. Importantly, most laminins exhibit high cell type specificity ^23^. The role of ECM components is well exemplified by recent work showing that the ECM skeleton of decellularized mouse heart can direct proliferation, differentiation, myofilament formation and electrical conductivity from human multipotent and pluripotent stem cell derived CMs to reconstitute a functional organ ^19,30^. However, which specific ECM components are responsible for these effects is, as yet, poorly understood.

Here, we show that LN-221 is the main laminin isoform expressed in cardiac muscle and we produced it for the first time as a human recombinant protein. When used as a culture coating for pluripotent hESCs, LN-221 induced a transcriptomic signature with upregulated markers for cardiac development as well as downregulation of tumor-inducing markers. Using LN-221 we have developed a fully chemically defined and xeno-free differentiation protocol. Importantly, this differentiation protocol is highly reproducible as demonstrated by remarkably similar transcriptomic profiles during differentiation of two different hESC lines to CM lineage for up to 94 days. We also identified the time points at which CVPs were generated, and further show with single-cell RNA sequencing (scRNA-seq) high reproducibility of the CVPs generated. We also identified three main subpopulations of cells with these CPVs namely cardiomyocyte-like cells (titin positive), fibroblast-like cells (lumican positive) and epithelial-like (epithelial cell adhesion molecule positive cells). Upon direct injection of 9- and 11-day differentiated CVPs into the heart tissue after pericardial injury, the CVPs differentiated further *in vivo* and formed cardiac muscle fiber bundles that survived in the heart for at least 12 weeks with improved cardiac function. These results suggest a role for laminins in CM differentiation and may allow the generation of clinical quality CVPs for regenerative cardiology in humans.

## RESULTS

### Generation of Recombinant Human Laminin LN-221 and Biological Significance for Cell Differentiation

In order to determine which laminins are most prominent in cardiac muscle tissue, we assessed whole transcriptome expression patterns of laminin genes in the human left heart ventricle of 108 non-diseased human donors ^31,32^. Analysis of the heart samples in this cohort showed that the *LAMB2* gene encoding the laminin *β*2 chain had the highest expression, followed by *LAMC1*, *LAMB1* and *LAMA2* that encode the laminin *γ*1, *β*1 and *α*2 chains, respectively (Fig. 1a). This identifies LN-221 as the most abundantly expressed laminin in the human heart. Ubiquitous BM laminins containing the *α*5 chain (*LAMA5*) and endothelial BM *α*4 chain (*LAMA4*) are also expressed in the left heart ventricle albeit at a lower level. We further investigated these genes at protein level and also observed *LAMA2*, *LAMB2* and *LAMC1* expression in human samples (Fig. S1a).

**Fig. 1.**
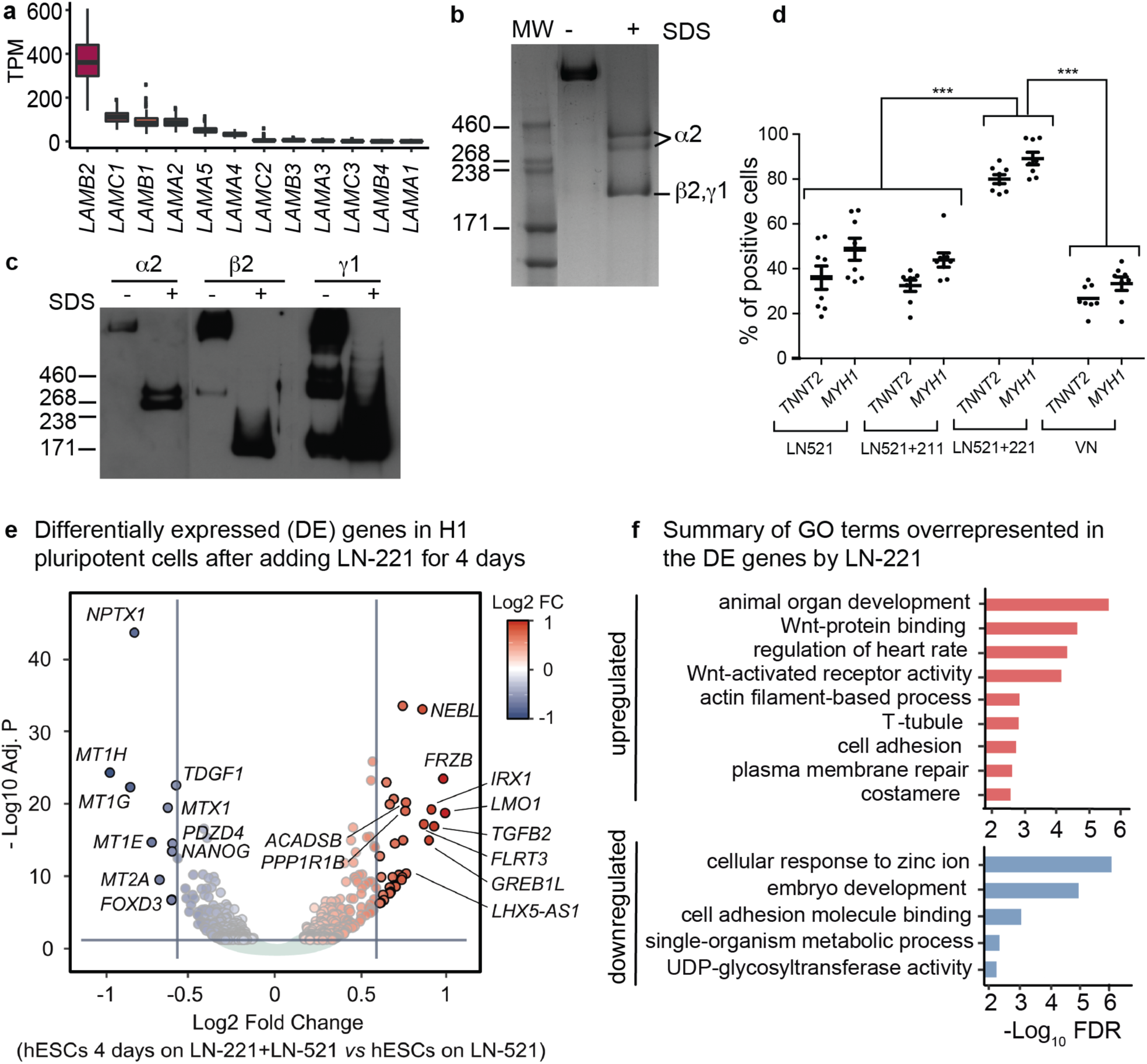
Generation of Recombinant Human Laminin LN-221 and Biological Significance for Cell Differentiation. **a** Laminin gene expression levels in heart left ventricle tissue retrieved from 108 non-disease human donors^31^. The most abundant laminin isoform was LN-221, which was used in the hESC to cardiomyocyte differentiation protocol developed in this study. See Methods for more details. **b** SYPRO Ruby red staining of molecular weight (MW) markers and recombinant human LN-221 analyzed by SDS-PAGE. The single band observed in non-reduced sample (-) demonstrates the purity of the laminin product. The two ~400 KDa bands observed after reduction (+) correspond to intact and partially processed *α*2 chains. The 200 KDa band seen in the reduced gel (+) corresponds to *β*2 and *γ*1 chains, which have similar molecular size. **c** Western blot image of individual laminin chains. LN-221 was produced in HEK293 cells as a disulfide-cross-linked heterotrimer. The *β*2*γ*1 dimer and monomeric *γ*1 chain can also be seen. (-) non-reduced, (+) reduced. **d** Combined flow cytometry measurements of *TNNT2* and *MYH1* in H1 cell. Using the combination of LN-521+221 significantly improved the efficiency of CMs production as compared to LN-521 alone, LN-521+211 and vitronectin (VN). *** P < 0.0001. **e** Initial transcriptomic experiment to inspect the cell differentiation potential of LN-221 in pluripotent cells. Volcano plot displaying the differential expression analysis of H1 hESCs cultured on LN-521 with addition of LN-221 for four days (n=3) comparing against H1 cells growing on LN-521 for four days (n=3). Gene names of the top ten up and downregulated genes (by fold change) are included. Genes are colored based on Log2 fold change (FC, X-axis). Genes with FC higher or lower than 1.5 are highlighted (±1.5 FC indicated by vertical lines). Y-axis: Benjamini–Hochberg (BH) adjusted p-values of the differential expression test (the horizontal line marks the 0.05 BH significance threshold, genes that did not pass this significance threshold are colored in light gray). **f** Summary of enriched Gene Ontology (GO) terms enriched in the set of up and downregulated genes (BH adjusted P<0.01), GO terms are summarized by semantic similarity). The full list of enriched functional terms is listed in Table S1. See Methods for more details.

Since LN-221 is difficult to isolate in pure form from muscle tissue, we generated it as a human recombinant in HEK293 cells, essentially as previously reported for other laminins LN-411 and LN-511^33,34^. Purified recombinant LN-221 is studied by SDS-PAGE after protein staining and by western blots (Fig. 1b, c). The native LN-221 did not penetrate into the gel under non-reducing conditions due to a disulfide-bonded heterotrimer conformation of over 700 KDa. However, after reduction three bands were observed, two bands of around 400 KDa corresponding to the intact *α*2 chain and its partially cleaved fragment and a 200 KDa band corresponding to *β*2 and *γ*1 chains, which have similar molecular size. The additional bands at lower molecular weight are presumably *β*2*γ*1 dimers but laminin dimers are not fully functional ^23^.

We have previously shown that recombinant human LN-521 alone supports pluripotency, stable karyotype, and expansion of hESCs *in vitr*o ^27^, which was confirmed here (Fig. S1b, S1c). In contrast, LN-221, when used as the sole culture coating, did not support attachment of hESCs. To explore the LN-221-induced effects on cardiac differentiation, using flow cytometry we compared the CM differentiation efficiency on four different matrices :LN-521 alone, LN-521 + LN-211, LN-521 + LN-221 and vitronectin (VN). The LN-521 + LN-221 matrix generated significantly more Troponin T (*TNNT2*) and myosin filament (*MYH1*) positive CMs than the matrix with LN-521 alone or LN-521 + LN-211 and VN (p<0.001) (Fig. 1d). LN-211 is closely related to LN-221, having an *α*1 chain in comparison with *α*2 in LN-221. LN-211 is also present in the BM surrounding individual CM fibers albeit in a lower amount (Fig. 1a). This strongly suggests that LN 521 + 221 is the most efficient matrix for cardiac differentiation.

In order to assess the potential of LN-221 to initiate and promote differentiation, we seeded single hESCs separately on a mixture of LN-521 + LN-221 and on LN-521 alone for four days and carried out bulk RNA-seq. We surmised that LN-521 would support the attachment required for cell proliferation, which in turn, would allow analysis of the effects of LN-221 on pluripotent cells. We observed that the LN-521 + LN-221 matrix indeed induced transcriptomic changes related to differentiation when comparing to the pure LN-521 matrix (Fig. 1e). Specifically, transfer of the pluripotent hESCs from day 0 to 4 on LN-221 led to upregulation and downregulation of 268 and 239 genes, respectively (false discovery rate (FDR) <0.05), which we denote as “LN-221 transcriptomic signature” (Table S1).

Genes with highest upregulation by LN-221 included *LMO1* (involved in neural and blood lineage development) ^35^, *IRX1* (role in embryonic patterning and congenital heart disease) ^36,37^, *TGFB2* (key regulator of heart development) ^38^, *FRZB* (involved in Wnt signaling) ^39^ and *NEBL* (involved in cardiac myofibril assembly) ^40^. Functional enrichment analysis (Fig. 1f, Table S1) of the upregulated genes revealed a main role in “animal organ development” (56 genes including *NEBL*, *KLF6* and *FZD3*, FDR=2.4 × 10^()^), “cell differentiation” (66 genes, among these *TGFB2*, *FGFR3* and *DSP*, FDR=7.72 × 10^()^), “Wnt protein binding” (7 genes including several members of the frizzled gene family, FDR=2.24 × 10^(+^) and “regulation of heart rate” (9 genes containing *TPM1*, *CACNA2D1* and *DMD*, FDR=4.45 × 10^(+^).

Interestingly, the ten most downregulated genes by LN-221 included tumor marker teratocarcinoma-derived growth factor *TDGF1* ^41^ and pluripotency marker *NANOG* ^42^. The downregulated genes also included the metallothionein-1H gene *MT1H* that is involved in “response to metal ions” (FDR=4.27 × 10^()^) and upregulated in both benign and malignant human tumors ^43^. Additionally, we found genes that are involved in “embryo development” (FDR=9.69 × 10^()^), such as *POU5F1* and neuronal marker *NPTX1* ^44^. In summary, these results suggest that LN-221-induced intracellular signaling initiates differentiation to the CM lineage and, importantly, seems to attenuate teratoma and tumor formation.

### Highly Reproducible Chemically Defined Xeno-free Protocol for Differentiation of hESCs to Cardiomyocytes

A key objective of this study was to investigate whether recombinant human LN-221 can initiate differentiation and partially support pluripotent hESC differentiation to the cardiovascular lineage. As a final goal, we aimed to develop a defined reproducible method for *in vitro* differentiation of hESC-derived CMs for the generation of clinical quality CVPs for treatment of heart injury. Therefore, no animal-derived matrix molecules were used in this study.

We developed a long-term differentiation protocol and collected samples for RNA-Seq of cells grown for 94 days from two separate cell lines hESCs lines, H1 and HS1001 ^27^ (Fig. 2a). The cells continued to differentiate into CVPs followed by CM formation without changing the matrix. Contraction of the cells was observed from day 13.

**Fig. 2.**
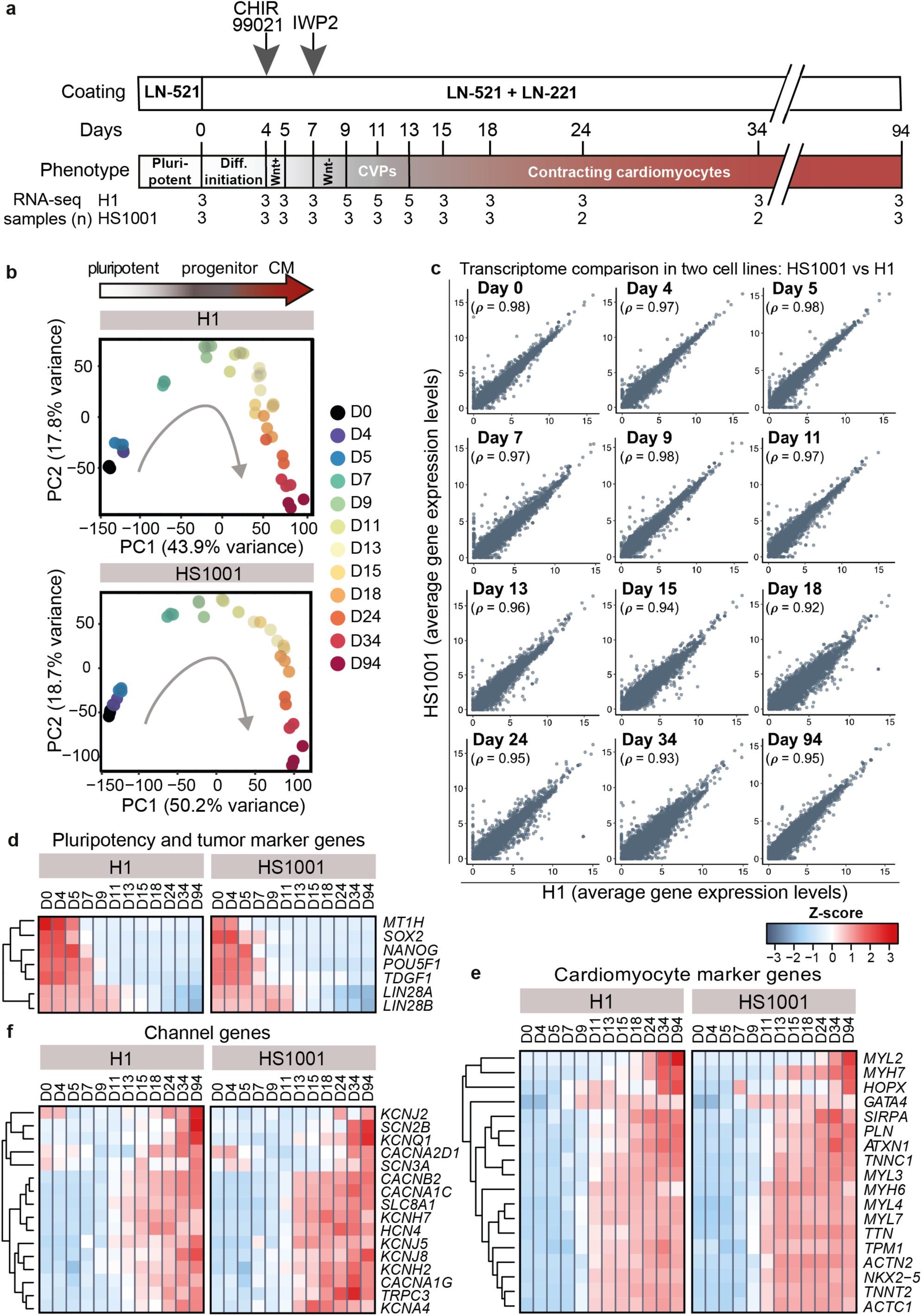
Highly Reproducible Chemically Defined Xeno-free Protocol for Differentiation of hESCs to Cardiomyocytes. **a** Schematic illustration of the LN-221 differentiation protocol. Pluripotent hESCs stably grown on LN-521 were trypsinized into suspension and re-seeded onto a mixture of LN-521 + LN-221 on day 0. Small molecule inhibitor CHIR99021 was added to the medium at day 4 to enhance Wnt signaling. On day 7, IWP2 was added to inhibit Wnt signaling. hESCs differentiated for up to 94 days were analyzed by RNA-seq for both H1 and HS1001 hESC lines. CMs differentiated for 34 days were subjected to detailed analyses as shown in Fig. 3. Two to five samples were studied by RNA-seq from the twelve different differentiation time points indicated at the bottom. **b** Principal component analysis (PCA) of pluripotent hESCs and cells collected throughout the differentiation protocol in H1 and HS1001 cell lines independently. Principal component (PC) 1 (x-axis) separates the data by time and PC2 (y-axis) captures cardiovascular progenitors (CVPs) formation. **c** Reproducibility of the differentiation protocol across time between H1 (X-axis) and HS1001 (Y-axis) cell lines. Whole-transcriptome expression levels correlation between H1 and HS1001 cell lines in all twelve time points. In the scatter plots, every point denotes a single gene (averaged across biological replicates in H1 and HS1001 respectively). *r* denotes Spearman’s rank correlation coefficient (rho). **d** Heatmap displaying the gene expression levels (Z-score of average transcripts per million, TPM) of teratocarcinoma/tumor (*TDGF1, MT1H*) and pluripotency genes (rest of genes) during the 94-day differentiation protocol in H1 and HS1001 cell lines. See Methods for details. **e** Heatmap with gene expression levels of mature CM markers during the 94-day protocol. (data are presented as in d) **f** Heatmap with gene expression levels of typical CM channels (data are presented as in d).

In order to assess reproducibility of the present differentiation method using LN-221, we profiled with bulk RNA-seq at twelve timepoints (from day 0 to 94), the transcriptome of cells differentiated with this protocol from two separate hESCs lines, H1 and HS1001 ^27^ (Fig. 2a). Principal component analysis (PCA) independently carried out on each cell line (Fig. 2b), revealed a time and possible CVP formation trajectory (captured by principal component 1 and 2 respectively). Importantly, this trajectory was highly similar between cells differentiated from both hESC lines. Comparison of the global gene expression profile of the two cell lines across all time points revealed exceptionally high reproducibility for all genes in the genome (Spearman’s rank correlation higher than 0.92 at all time points, Fig. 2c). Time expression changes of known marker genes for pluripotency and tumor markers (Fig. 2d), CMs (Fig. 2e) and cardiac channel genes (Fig. 2f) exhibited alike transcriptomic expression profiles between the two cell lines. This high reproducibility suggests that the present differentiation protocol may yield cells that can be safe for use in therapy. Based on the expression levels of multiple mature CM markers (e.g. *TNNT2*, *ACTC1*, *MYH7* and *SIRPA* (Fig. 2e), differentiation to contracting CMs takes place around day 13. This was also confirmed by the development of a beating cellular phenotype (Supplementary information, Movie S1) and microscopy and immunohistological analyses at day 34 (Fig. 3).

**Fig. 3.**
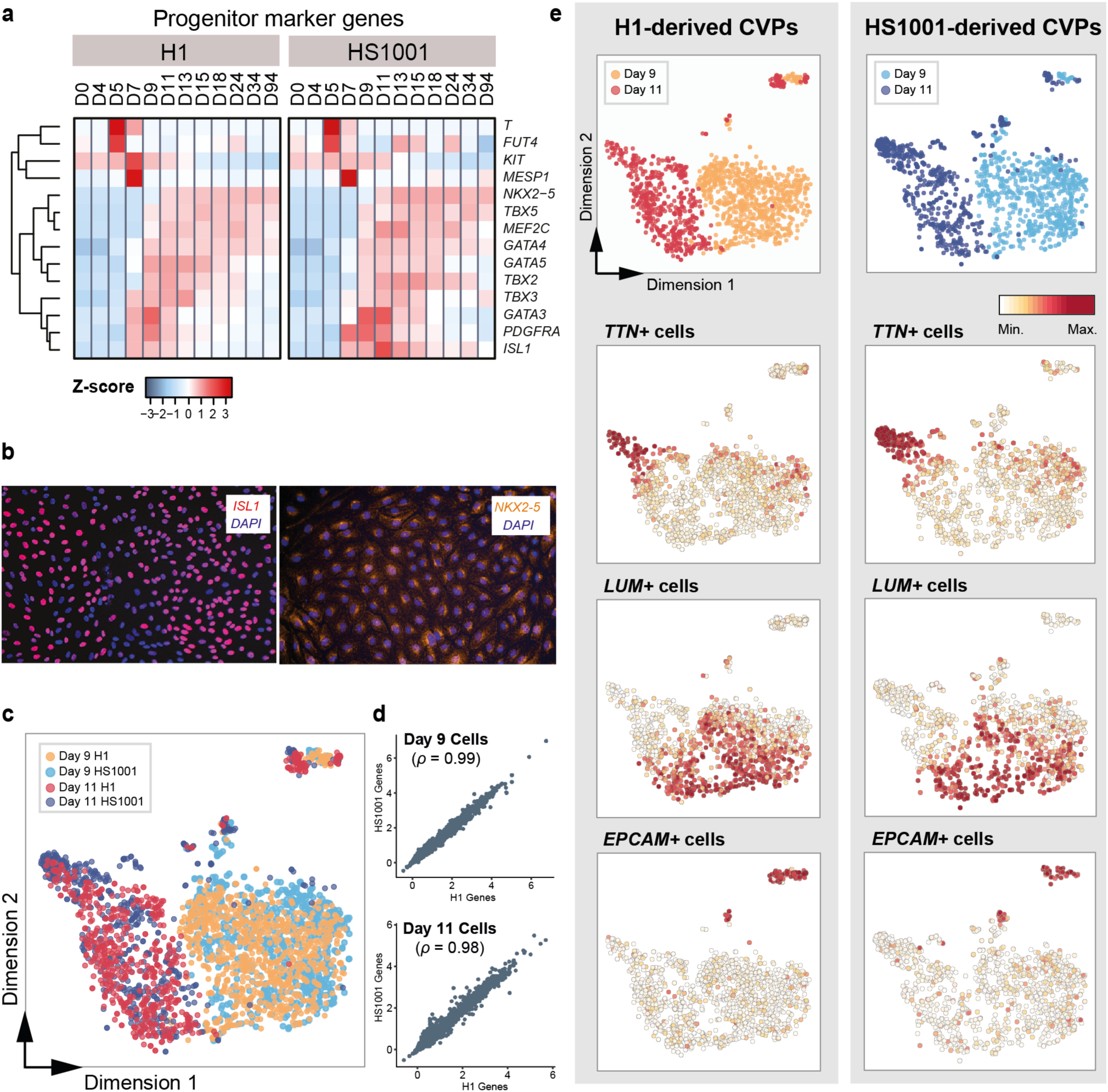
Identification and Characterization of CVPs. **a** Heatmap showing the gene expression levels (average TPM Z-scored) of known progenitor markers across the differentiation time-course in both H1 and HS1001. **b** Left, H1-derived CVPs at day 9 of differentiation stained with ISL1 antibody (Red) and DAPI (Blue). Right, H1-derived CVPs at day 9 of differentiation stained with NKX2-5 (Orange) and DAPI (Blue). **c** Graph showing the first two t-SNE components computed from all four single-cell sequencing runs together (day 9 and 11 H1- and HS1001-derived CVPs). Each dot denotes an individual cell. Cells are shown mapped to condition (i.e. day and hESC line). **d**. Scatterplot showing the correlation level of all genes across all cells profiled in the two cell lines for day 9 (top) and 11 (bottom) CVPs. In these graphs, each dot denotes a the average expression (logged counts after accounting for cell cycle effects) of a single gene across all cells. *r* denotes Spearman’s ranked correlation. **e** Same graph as in C but now only showing only either H1- or HS1001-derived CVPs, left and right respectively. The top graphs display the cells coloured by time point whereas the graphs below display each cell mapped to the expression level of genes marking three main subpopulation of cells: Titin (*TTN*), Lumican (*LUM*) and Epithelial Cellular Adhesion Molecule (*EPCAM*).

### Characterization of hESC-derived Cardiomyocytes after 34-Day Differentiation *in Vitro*

From the CM generated using this protocol, we observed unidirectional alignment of contracting cells mimicking muscle fibers and/or whole sheet of beating CMs (Supplementary information, Movie S1). The cell sheets were dissociated and single CM were re-plated onto plates coated with LN-521 + 221. The CMs attached and spread on the coating forming spindle-shaped, rectangular and cuboidal-shaped morphologies resembling isolated adult CMs ^45^ (Fig. S2a). Immunofluorescence staining with heart muscle specific Troponin I (*TNNI3*), *TNNT2* and *α*-actinin antibodies clearly showed sarcomeric striations, as well as the presence of bi-nucleated cells with *NKX2-5* staining (Fig. S2b-d) which implied a CM phenotype. Cardiac markers were also probed with specific antibodies against *TNNT2* and *MYH1* in flow cytometry analysis and we were able to generate >80% positive CMs from both H1 (Fig. S2e-f) and HS1001 cell lines (Fig. S2g-h).

To further characterize the functionality of the cells, single cell patch clamp and multi-electrode assay (MEA) were carried out to measure the CMs action potential and responses to drug compounds. The CMs matured for 32-40 days exhibited significant similarities in electrophysiological properties as previously reported for hPSC-derived CMs ^46–48^. Ventricular-, atrial- and nodal-like CMs were identified with proportions of 64.2%, 22.2% and 13.6%, respectively (Fig. S2i, Table S2). Functionality and relevant responses to pharmacologic compounds were demonstrated in a cell viability assay (Fig. S2j-k). Treatment with valinomycin or emetine dihydrochloride showed that the drugs significantly reduced cell viability with various potencies. The EC_50_ values calculated were 13.1 μM for valinomycin and 6.079 μM for emetine dihydrochloride, which demonstrated the sensitivity of the cells to cytotoxic compounds. Cellular responses to adrenergic stimulation were also tested with multi-electrode array. Upon exposure to isoproterenol and sotalol, the CMs increased heart rate (Fig. S2l) and shortened corrected field potential duration (cFPD), respectively (Fig. S2m and Table S1). These results agree with previously published studies in which they also found induced pluripotent stem cells derived CM responded equally to isoproterenol^49^ and sotalol ^50^. In conclusion, *in vitro* differentiation of hESC to beating CMs with our laminin-based, defined and xeno-free protocol yielded reproducibly CMs with properties highly similar to those isolated from intact heart muscle.

### Identification and Characterization of CVPs

CVPs have been reported to have high plasticity and be able to differentiate into different several heart cell types so they would be useful for transplantation into infarcted hearts which may potentially avoid the problem of arrhythmia ^51^. The rationale for using 9- or 11-day differentiated CVPs was that these cells have low expression of pluripotency and tumorigenicity genes (Fig. 2d), low expression of CM genes (Fig. 2e), high expression of known CVPs genes, such as *ISL1*, *C-KIT* and *GATA3* (Fig. 3a) and they still proliferated efficiently *in vitro*. Moreover, the decline in pluripotency and tumor markers highly suggested that the cells have low tumorigenicity and might be safe for transplantation.

To identify the timepoint(s) at which the cells reach progenitor stage with our protocol, we inspected the time series transcriptomic profile of known progenitor marker genes (Fig. 3a). In cells derived from both H1 and HS1001 hESCs, we found a significant strong upregulation of the mesendoderm marker Brachbury gene (*T*) at day 5. We detected further specification into mesoderm lineage at day 7, marked by significant upregulation of *MESP1* ^52^. *ISL1* gene has been previously shown to mark early cardiovascular progenitor stage ^53^. In our cells, we observed significant upregulation of *ISL1* from day 7. However, other CVPs markers such as *TBX5*, and *MEF2C* did not show significant upregulation until day 9. In day 9 H1-derived cells, we also confirmed the protein expression of ISL1 and NKX2-5 (Fig. 3b). NKX2-5 gene has been reported to mark early CM and we observed its most significant upregulation at day 11, suggesting that cells at day 11 may also capture early cardiomyocyte-like features. Therefore, we defined our CVPs as day 9 and 11 cells.

To further assess the reproducibility of our protocol and characterize the heterogeneity of the CVPs, we carried out scRNA-seq in H1- and HS1001-derived CVPs from day 9 and 11. The t-Distributed Stochastic Neighbor Embedding (t-SNE) visualization uncovered an overlapping profile for CVPs derived from the two independent hESC lines (Fig. 3c). Whole transcriptome correlation analysis revealed a very high correlation between the transcriptome of CVPs derived from H1 and HS1001 at both day 9 and 11 (Fig. 3d, Spearman’s ranked correlation level of 0.99 and 0.98 for day 9 and 11 cells respectively). This confirmed the very high reproducibility observed across time as observed previously with bulk RNA-seq (Fig. 2c) and further emphasizes the high reproducibility of the laminin differentiation method. We also inspected the cellular heterogeneity present in these CVPs and in H1- and HS1001-derived CVPs from day 9 and 11 we uncovered three main sub-populations of cells marked by the expression of the genes titin (*TTN*), luminan (*LUM*) and epithelial cell adhesion molecule (*EPCAM*) (Fig. 3e). Although the cells expressing EPCAM displayed marked differences, the rest of the cells showed an opposing gradient towards either higher lumican or titin expression.

### Transplanted Progenitor Cells Form Striated Human Cardiac Muscle Fibrils and Neovascularization in Infarcted Region

To determine whether the CVPs can generate new heart muscle tissue *in vivo*, we used a cardiac ischemia/reperfusion (I/R) injury model in mice and administered luciferase-labeled CVPs to damaged myocardium. No mortality of mice occurred during the follow-up and the study design is summarized in Fig. S3a. Flow cytometry analysis confirmed that luciferase expressing lentivirus-transfected hESCs maintained their pluripotency from the expression of Tra1-60 and *POU5F1*. (Fig. S3b).

Safety concerns regarding stem cell-derived neoplasia were addressed using whole body IVIS imaging. Luciferase signal was only detectable in the heart but not in other organs suggesting non-invasive nature of the progenitors and long-term viability throughout the 12-week follow-up (Fig. S3c). Teratomas were not observed in any of the progenitor-treated nude mice after 12 weeks, which strongly suggests that the cells were not immunogenic and are likely to be safe for cell therapy (Fig. S3d-e).

We first assess the reproducibility and stability of our surgical method, we measured the area at risk (AAR) and infarct size of the ischemic tissue 24 hours after coronary occlusion (Fig. S3f-k). We observed no differences in any of the groups (medium- or progenitor-treated) suggesting the high reproducibility of our surgical technique.

Mouse hearts were excised for histological analysis at 12 weeks post-transplantation. Photos of native hearts at the time of sacrifice demonstrated scar formation (outlined in yellow) and absence of teratoma development (Fig. 4a). This was evident from heart sections stained with Picrosirius Red dye to display presence and extent of fibrotic lesions (Fig. 4b). Hearts treated with medium alone had larger scar compared to the hearts treated with progenitor cells (p<0.05) (Fig. S3l). We observed a reduction in scar volume in hearts treated with 9–day progenitors as compared to medium-treatment, but this was not significant (p = 0.054). We also observed thinning of the ventricular walls in medium-treated hearts suggesting the adverse ventricular remodeling (Fig. 4b).

**Fig. 4.**
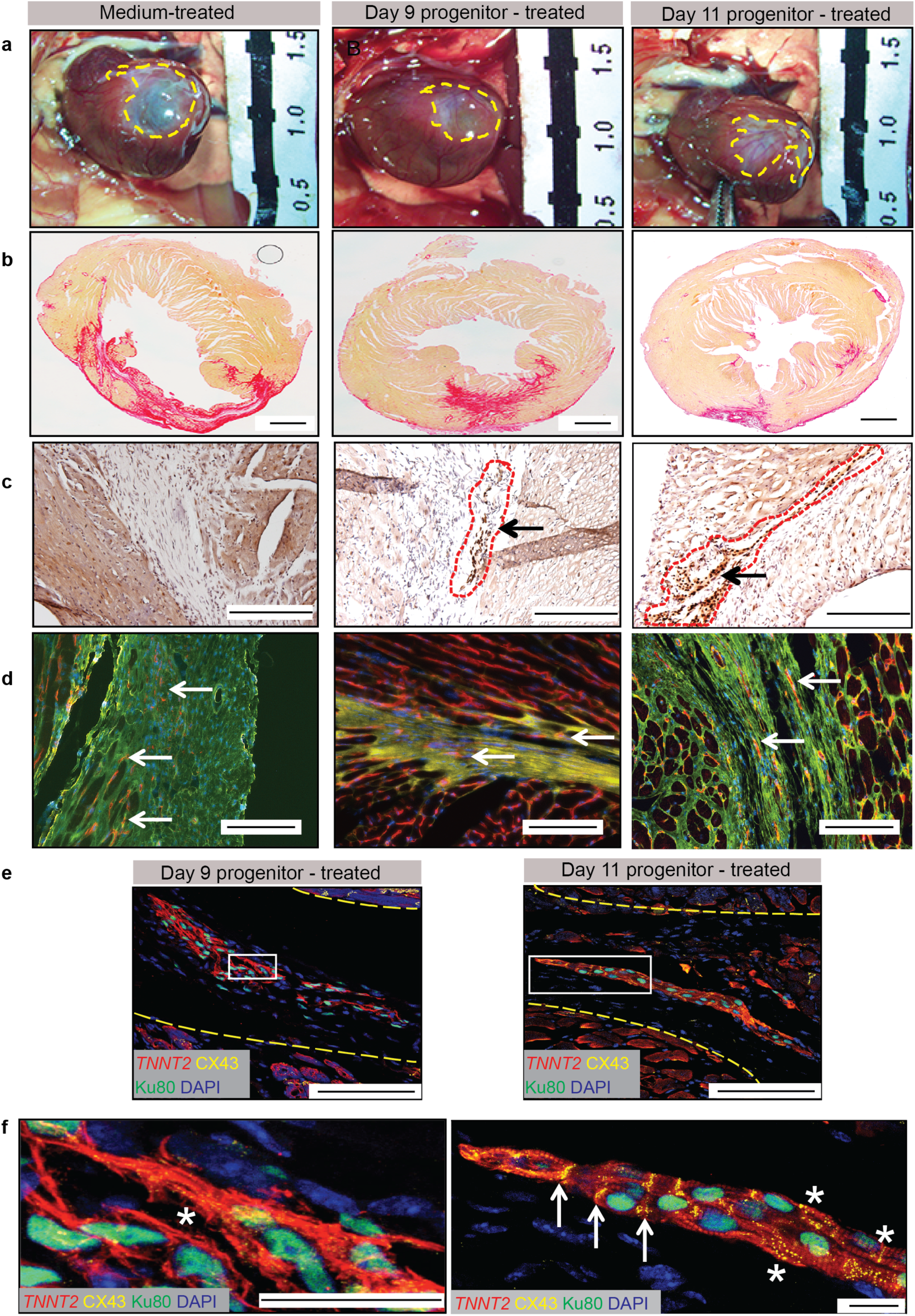
Transplanted Progenitor Cells Form Striated Human Cardiac Muscle Fibrils and Neovascularization in Infarcted Region. **a** Photomicrograph of mouse heart at the time of sacrifice (12 weeks post MI). The yellow outline demarcated the region of infarction. **b** Picrosirius Red staining reveals that the fibrotic area was larger in hearts treated with medium only than in progenitor-treated hearts. Fibrotic tissue is stained red and viable tissue remains pink. There is significantly lower scar volume (p<0.05) in day 11 progenitor-treated as compared to medium-treated (Refer to Fig. S3l). Scale bar = 1 mM. **c** 3,3’-Diaminobenzidine staining for nucleic acids (brown) together with human nuclei specific Ku80 antibody shows areas in the infarcted region containing human cells. Positive human cells were outlined in red. Scale bar = 200 μM. **d** Angiogenesis determination using isolectin B4 (red) and wheat germ agglutinin (WGA, green) antibody. There are significantly more neovessels (red) (P<0.05) in day 9 progenitor-treated hearts as compared to medium-treated (also refer to Fig. S3k). Scale bar = 100 μM. **e** Confocal images showing the presence of human muscle fiber bundles double-stained stained with *TNNT2* (red), human nuclei Ku80 (green) and connexin 43 (Cx43, yellow) at 40X magnification. Scale bar = 25 μM. **f** Confocal images of muscle bundles in the white box in panel (e) at 100X magnification. Scale bar = 25 μM. The yellow dotted line demarcates the region of infarction. Well-organized and parallel cytoplasmic striations are observed in hearts treated both with progenitors differentiated for both 9 and 11 days. White arrows point to areas positive for Cx43 the end plates between two CMs, and asterisks indicate areas with less organized Cx43 protein. The presence of Cx43 clearly identifies the end-to-end connections between CMs in the muscle fiber bundles.

Immunostaining of the I/R injury region with an antibody specific for human nuclei revealed clusters of human cells (encircled by a red dotted line in Fig. 4c). Neo-angiogenesis was revealed in the I/R injury area by staining blood vessels with isolectin B4 (red) and fibrotic area stained with anti-wheat germ agglutinin (green) (Fig. 4d). Significant increase in the number of blood vessels was observed in the I/R injury region of day 9 progenitor-treated hearts as compared to solely medium-treated hearts (p < 0.05) (Fig. S3m).

The presence of multiple human nuclei was observed in cardiac troponin T (*TNNT2*) positive muscle fiber bundles in the mouse I/R injury region using an anti-human DNA helicase-specific Ku80 antibody (Fig. 4e). At 100X magnification, *TNNT2* striations and human DNA helicase staining in green were readily seen (Fig. 4f). This demonstrates that the transplanted troponin T negative CVPs matured to CMs *in vivo*. Furthermore, an antibody against connexin 43 (Cx43) (yellow) highlighted end-to-end gap junctions that are essential for the formation of functional CM bundles (Fig. 4f, Supplementary information, Movie S2). Cardiomyocytes generated by “11-day progenitors” formed well-organized bundles of muscle fibers containing intercalated discs typical for cardiac muscle. The intercalated discs generated from “9-day progenitors” had a more dispersed arrangement at one end suggesting an ongoing aggregation and maturation of muscle tissue. Overall, the presence of Cx43 demonstrates that CVP-derived human muscle fibers are well connected to each other and are most likely functional.

### Transplantation of Cardiovascular Progenitors into Acutely Infarcted Mouse Ventricle Restores Myocardium Contractile Function and Attenuates Left Ventricle Deformation and Dyssynchrony

The cardiac function after I/R injury was investigated using echocardiography. As expected, left ventricular impairment and remodeling was observed in all three experimental groups after I/R injury (Fig. 5a and Table 1). The anterior wall was dramatically impaired in the medium-treated controls as compared to the baseline control mice at week 8 post-surgery (Fig. 5a). However, this impairment was significantly attenuated in the mice treated with CVPs, which also had thicker walls (Fig. 5a).

**Fig. 5.**
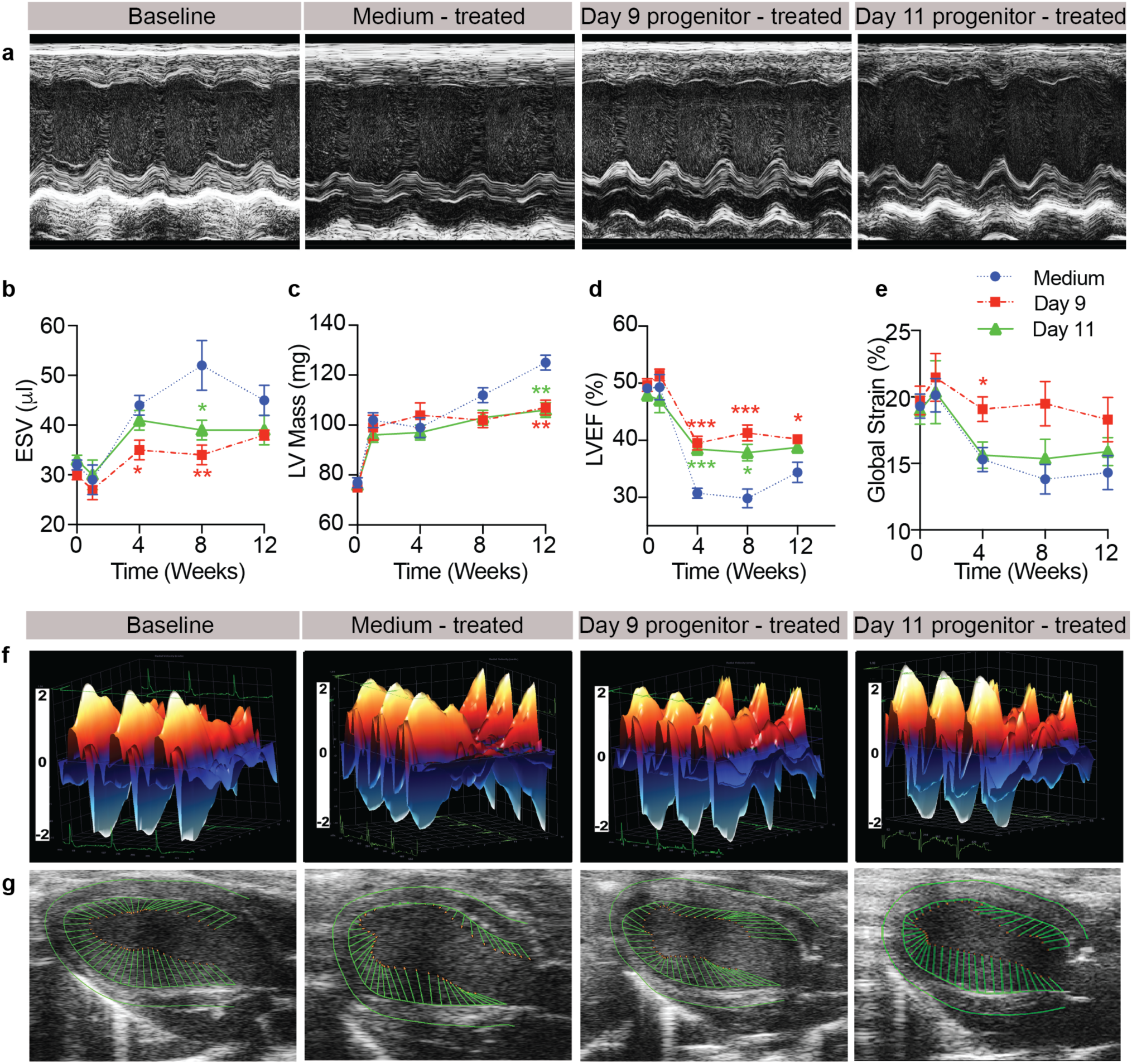
Transplantation of Cardiovascular Progenitors into Acutely Infarcted Mouse Ventricle Restores Myocardium Contractile Function, Attenuate left Ventricle Deformation and Dyssynchrony. **a** Representative echocardiogram (M-mode tracing) of left ventricle (LV) at baseline, medium and progenitor-treated mice. **b** End systolic volume (ESV), **c** LV mass and **d** left ventricle ejection fraction (LVEF) measured by echocardiography, and **e** radial strain measured by speckle-tracking strain analysis. Data were presented as Mean ± SEM and analyzed by 2-way ANOVA with Bonferroni *post hoc* analysis. *p < 0.05; ** p < 0.01; *** p < 0.001 n = 12 for medium control; n = 9 for day 9 CVPs and n = 7 for day 11 CVPs. **f** 3D regional wall velocity diagrams of LV endomyocardial strain showing contraction (orange) and relaxation (blue) of 3 consecutive cardiac cycles. **g** Vector diagrams showing direction and magnitude of endocardial contraction at mid-systole.

Next, we measured heart chamber volumes at baseline (week 0) and weeks 1, 4, 8 and 12 (Fig. 5b). We observed that in medium-treated mice there was a significant increase in end systolic volume (ESV) at 4 to 8 weeks after myocardial I/R injury (p<0.05) (Fig. 5b and Table 1). In contrast, increase in ESV was attenuated at week 4 (for day 9 progenitor-treated) and week 8 (for day 9 and day 11 progenitor-treated) in the progenitors-treated mice. The changes in end diastolic volume (EDV) were relatively moderate and the attenuation in EDV reached statistical significance (p<0.05) only at week 8 for mice treated with day 9 progenitors compared to medium-treated mice (Table 1). Eccentric cardiac remodeling (as indicated by the increase of left ventricle mass in LV mass) was induced following myocardial I/R injury in all mice (Fig. 5c). Compared to the medium-treated mice, the mice treated with CVPs showed significantly less remodeling (p<0.01) at week 12 after myocardial I/R injury (Fig. 5c).

Cardiac function, as indicated by left ventricular ejection fraction (LVEF), was reduced by myocardial I/R injury (Fig. 5d and Table 1). Compared to medium-treated mice, transplantation of day 9 progenitors markedly improved cardiac function by ~ 30 % (p<0.001). Similar treatment efficacy was achieved with day 11 progenitors at weeks 4 and 8. Similar to LVEF, fraction shortening (FS) was dramatically decreased (p<0.01) from week 4 onward in medium-treated mice and the decrease was alleviated by transplantation of day 9 or 11 progenitors (Table 1). In order to assess the left ventricular deformation, we measured the global strains parameters. Treatment with day 9 progenitors significantly (p<0.05) alleviated the decline in global radial strain after myocardial I/R injury at week 4 (Fig. 5e). However, we did not observe statistical global radial strain improvement after the transplantation with day 11 progenitors.

To investigate whether transplantation of CVPs can improve contractile activity in the I/R injured region (also the cell injected region), speckle-tracking strain analysis was performed with three consecutive cardiac cycles. Speckle tracking strain analysis is an imaging technique that analyses the velocity at which deformation of the tissues occurs in the heart in comparison to the tissue’s original dimension. To depict treatment efficacy of cell transplantation, representative data obtained at week 8 after myocardial I/R injury are presented in Fig. 5f-g. As demonstrated by the three-dimensional (3D) wall velocity diagrams, there was a marked reduction in wall velocity across the anterior wall (I/R injured region for cell transplantation) in medium-treated mice (Fig. 5f). The anterior wall velocity showed improvement in mice treated with day 9 or day 11 progenitors (Fig. 5f). Subsequently, we investigated heart function using vector diagrams and live echocardiogram video loops for all the time points (Fig. 5g). At baseline, the heart shows synchronous contractions and relaxations throughout the cardiac cycles (Fig. 5g and Supplementary information, Movie S3). Eight weeks after myocardial I/R injury, mice treated with medium only showed overt chamber dilation and hypokinesis of the anterior wall as evidenced by the reduction of vector activity in that region (Fig. 5g and Supplementary information, Movie S4). In contrast, the contractile activity of the anterior wall was significantly improved in mice treated with day 9 progenitors (Fig. 5g and Supplementary information, Movie S5). Improvement in contractile activity of the anterior wall in mice administered with day 11 progenitors was also observed although to a smaller extent (Fig. 5g and Supplementary information, Movie S6). Taken together, these results demonstrate that the treatment with CVPs restored heart function and attenuated LV deformation after myocardial I/R injury.

## DISCUSSION

The present results describe a highly reproducible laminin-221 based xeno-free protocol for differentiation of pluripotent hESCs to well-defined CVPs that, when injected into acutely infarcted region of mouse heart, proliferate and mature to form bundles of human cardiac muscle fibers. We consider these results important for several reasons.

First of all, we showed that the biologically relevant laminin isoform LN-221 is the most abundantly expressed BM laminin in the heart and that initiate differentiation of pluripotent hESCs to CMs and maintain reproducible differentiation. It was produced in this study as a human recombinant protein and can be produced under Good Manufacturing Practice (GMP) conditions for clinical applications. The addition of LN-221 to a LN-521 culture coating, induced transcriptomic signatures to increase Wnt signaling in hESCs. Intriguingly, during the 94-day differentiation protocol we observed highly similar global expression profiles in cells grown from two hESC lines generated by different laboratories, methodologies and even decades apart ^27,54^. This emphasizes the reproducibility of the method and the importance of using pure biologically relevant components in the cell culture matrix that can mimic the *in vivo* cell niche. Secondly, our protocol is fully xeno-free and chemically defined, which may allow this method to be used for production of cells for clinical regenerative cardiology. Thirdly, we did not find any evidence for tumorigenicity in the CVPs used in pericardial injections based on whole body imaging and histological analyses. Finally, the transplanted cells formed human muscle fiber bundles and improved heart function in mice with cardiac I/R injury. Therefore, our differentiation method may fulfil the requirements of regulatory authorities for generation of therapeutic stem cell derived cells for regenerative cardiology.

This is the first time that LN-221 has been produced, explored and shown to impact hESC differentiation. While LN-521 supports growth of pluripotent hESCs ^27^, plating of the pluripotent cells on a LN-521 + LN-221 matrix for four days initiates differentiation. LN-221 is likely to play a role in the development of cardiac muscle *in vivo*, although addition of “pharmacological” amounts of CHIR99021 was found to be necessary for boosting the differentiation *in vitro,* as previously published ^55^. The use of the combined LN-521 + LN-221 matrix gives high reproducibility and stable cardiovascular differentiation. The identification of a “LN-221 transcriptomic signature” reveals a mechanistic insight into how this laminin promotes differentiation towards CVPs and CMs.

There are efficient methods for differentiating hESCs to CMs ^56,57^ but they have involved the use of complex, undefined constituents that form an obstacle for the use of such cells in the clinic, and therefore are not appropriate for making therapeutic cells. One study reported a “chemically defined” protocol for the generation of human CMs using a non-biologically relevant vitronectin molecule and Matrigel^TM^ ^56^. This protocol was xenogenic and undefined, and it did not generate cells that were functional in an animal model. However, in agreement with our studies, these investigators also reported higher growth-rates and long-term CM adhesion on a LN matrix, as we have reported previously ^27^. There are other laminin-based CM differentiation protocols (LN-521^58^, LN-511^56^, LN-211^59^) to generate CM from pluripotent stem cells with similar differentiation efficiency, nevertheless these studies did not investigate the biological effects of laminins on the differentiation and did not carry out transplantation of differentiated cells to repair injured heart.

Injection of 9- or 11-day differentiated CVPs resulted in a significant improvement in cardiac function with survival of the administered human cells and formation of muscle fiber bundles containing well-formed intercalated discs. This may be partially due to the formation of angiogenesis that is necessary for the growth of new tissue as observed from histology. Injection with day 11 CVPs resulted in a better arrangement of the muscle fibers as compared with day 9 CVPs. This could be due to higher expression of cardiomyocyte-like genes at day 11 CVPs as shown by both our bulk RNA-sequencing and scRNA-sequencing. Interestingly, the ECM proteoclycan agrin has been shown to promote heart regeneration in mice by facilitating CM cell cycle reentry ^60^. Agrin is found in most membranes and, thus, does not exhibit the same cell type specificity as the laminins do. It would be interesting to explore whether a combination of agrin and our CVPs can further enhance cardiac performance. There is also a recent study by Zhu *et al* that reported the lack of remuscularization of the nonhuman primate hearts after transplantation of SSEA-1 positive (*FUT4*) CVPs ^61^. It is important to note that their cells had only been differentiated for 3 days and when injected into infarcted hearts, they could not survive long-term. In our RNA sequencing data, we observe the highest expression of *FUT4* gene on day 5, but its expression is very low in 9- and 11-day differentiated CVPs used in our study (Fig. 3a). We postulate that Zhu *et al* used cells that were at a too early differentiation stage and not capable of remuscularization. Hence, the present work suggests that our CVPs are appropriate cells to be used for cardiac regeneration.

For future work, our scRNA-seq analysis of CVPs uncovered three main sub-population of cells at days 9 and 11. It will be important to separate and sort these sub-populations and explore if any of these can enrich for cell types that intensify the growth of new heart muscle *in vivo*.

## MATERIALS AND METHODS

### Laminin Expression of the Left Ventricle Human Heart Tissues

To select the laminin to be used in the differentiation protocol, we analyzed RNA sequencing (RNA-Seq) data from non-diseased human heart left ventricle. We analyzed the 108 non-diseased controls previously published by Heinig *et al* ^31^.

Transcripts Per Million (TPM) were computed from gene counts as following: gene read counts were divided by gene length in kilobases to obtain reads per kilobase (RPK, gene length was computed by summing up the length of the total number of exons in each gene using the R library GenomicFeatures 1.26.3) ^62^. To obtain TPM values, these RPKs were divided by the sample “per million” scaling factor (defined as the total number of gene counts in a sample divided by a million). Data was quantile normalized by using the function *normalize.quantiles* from R package preprocessCore 1.36.0 ^37^. All laminin genes were selected and TPM levels were visualized in a boxplot (Fig. 1a).

### Production and Characterization of Human Recombinant Laminin-221 (LN-221)

The LAMA2 cDNA (NCBI gene ID: 3908) was synthesized and cloned into expression vector pcDNA3.1/Zeo (-) (Invitrogen). DNA sequencing was employed to ensure the complete cloning of LAMA2 (9396bp) cDNA in the correct reading frame and direction. Constructs used for the expression of full-length human *β*2 and *γ*1 chains have been described previously ^27,33^. To produce the full-length trimeric LN-221 (BioLamina AB, Sweden), individual plasmids containing *α*2, *β*2 and *γ*1 chains were sequentially transfected into human embryonic kidney cells (HEK293, ATCC CRL-1573) as previously described for the production of LN-511 ^34^.

Purified LN-221 was characterized using gradient SDS-PAGE. Proteins were visualized using SYPRO Ruby staining (Bio-Rad) or transferred onto PVDF membranes (Bio-rad). The membranes were probed with polyclonal antibodies against respective chains (anti-*α*2, C13065, anti-*β*2, CC13070 and anti-*γ*1, CC13072, AssaybioTech). After washing, the membranes were incubated with horseradish peroxidase-conjugated donkey anti-rabbit IgG (1:5000). Chemiluminescence kit (GE Healthcare) was utilized to detect the immunoreactivity as according to the manufacturer’s instructions.

### Analysis of LN-221 Bulk RNA-Seq Data to Assess the Effect of LN-221 on Pluripotent Stem Cells

Differential expression analysis was carried out with DESeq2 1.14.1^63^. DESeq2 was run pairwise (LN-521+LN-221 samples were compared to LN-521 samples using Wald test). In DESeq2 *results* function, the *independentFiltering* parameter was set to True, the rest of parameters were left as default. Genes were considered significantly differentially expressed (DE) if Benjamini & Hochberg (BH) adjusted p-value < 0.05.

Functional enrichment of the DE gene sets was computed using the function *gprofiler* from R package gProfileR version 0.6.1 ^64^. We tested two DE gene sets: genes with BH adjusted p-value < 0.05 split into positive and negative fold change. The background was set to the input set of genes in DESeq2. External gene names of the DE genes were used as query. Electronic annotations were excluded, the p-value correction method was set to “fdr” and only results with FDR smaller than 0.05 were considered. For visualization purposes, summary of results from GO enrichments (BP, MF and CC) were generated by filtering terms with a high semantic similarity ^65^, using the Lin distance measure with a threshold of 0.3. The obtained list of summarized GO terms (FDR<0.01) was visualized in a graph (Fig. 1f). See Table S for the full list of enriched terms in the up and downregulated genes.

### Generation of Bulk RNA-Seq Data

Three separate bulk RNA-Seq experiments were carried out. We performed one initial experiment to assess the effect of LN-221. In this experiment, we sequenced pluripotent H1 cells growing on LN-521 and LN-521+LN-221 for four days (n=3 each). We also carried out two time-series experiments with cell differentiated from cell lines H1 and HS1001 at different days from the differentiation protocol (at day 0 (pluripotent stage), and days 4, 5, 7, 9, 11, 13, 15, 18, 24, 34 and 94). The number of biological replicates sequenced in both H1 and HS1001 cell lines can be seen in Fig. 2a. HS1001 cell line was sequenced with three biological replicates at each time point (apart from day 24 and 34 with two replicates). We denote these three experiments as LN-221 dataset, H1 time-course, HS1001 time-course.

In all three experiments, RNA was isolated from the CMs at specific time points with single-cell RNA purification kit (Norgen Biotek Corporation) according to manufacturer’s guidelines. RNA quality was measured with bioanalyzer (Agilent) and RNA with RIN number at least 8 were used for RNA-Seq at Duke-NUS Genome Biology Facility using Illumina Hiseq 3000 RNA-Seq machine. Libraries were constructed with TruSeq stranded total RNA library Prep Kit (Illumina). In the two time series runs samples were multiplexed with 10 samples per lane. All six samples were run in a single lane in the case of the LN-221 dataset. Each of the three experiments was run in a separate sequencing batch. These three RNA-seq datasets have been deposited in NCBI’s Gene Expression Omnibus ^66^ and are accessible through GEO Series accession number GSE100725.

Reads were assessed for quality and (when necessary) adaptors were removed using Cutadapt ^67^. RNA-Seq reads were aligned to hg38 (Ensembl Gene annotation build 79) using STAR 2.5.2b ^68^ and quantified using RSEM 1.2.31 ^69^, with an average mapping rate of 97.9% for H1 time-course, 98.2% for HS1001 time-course and 95.6% for LN-221 dataset. Gene annotation was retrieved from Ensembl version 79 (hg38) using the R library biomaRt 2.30.0 ^41^. Ribosomal genes (Ensembl gene biotype “rRNA”) and mitochondrial genes were removed (584 genes in total). The highest expressed outlier small non-coding RNA genes were also removed (“RN7SL1” and “RN7SL2” genes, in the LN-221 dataset “RN7SK” was also removed). Gene counts were rounded using the R function *round* ^70^. A pre-filtering step was added in each dataset, considering only genes with more than 1 count when summing up across all samples.

### Analysis of the Reproducibility of the Differentiation Protocol Using H1 and HS1001 Bulk RNA-Seq Time Series Data

We assessed the reproducibility of the differentiation protocol using the two time-course bulk RNA-Seq datasets (H1 and HS1001 cell lines). We correlated genome-wide average TPM levels between both cell lines across each time point (Fig. 2b). In addition, we used principal component analysis (PCA) to inspect patterns when reducing the dimensionality of the data. For this PCA analysis, a *DESeqDataSet* object was created with the design formula (~RIN score + time variable). Regularized-logarithm transformation was subsequently applied using the function *rlog* (DESeq2 1.14.1) ^63^ and setting the parameter *blind* to false. PCA was computed with the R function *prcomp* with parameters *center* set to true and *scale* set to false ^70^. The percentage of contribution of each gene to PC1 and PC2 was also computed.

Expression levels of pluripotency, CM, channel and cardiovascular progenitor genes were visualized throughout the protocol. Log2(TPM+1) in H1 and HS1001 were computed. Then, the data was combined into a single matrix and it was quantile normalized using the function *normalize.quantiles* from R package preprocessCore 1.36.0 ^37^. Z-score of all genes was computed in the joint dataset. Each of these set of genes were visualized clustered according to the mean Z-score in H1 cell line using the function *heatmap.2* from R package gplots 3.0.1^13^. Pluripotency markers were retrieved from Takahashi et al ^71^ and CM markers and channels genes retrieved from ^72^. In addition, differential expression analysis was run with DESeq2 1.14.1 ^63^ both pairwise comparing all time points and also considering all time points together. RNA quality score (RIN) was added as a covariate in the model. In DESeq2 results function, the *independentFiltering* parameter was set to False, rest of parameters were set as default. Wald tests were carried out in the case of pairwise comparisons and likelihood ratio test (LRT) was used to test for genes varying across all time points (including pluripotent cells). Genes were considered significantly differentially expressed (DE) if BH adjusted p-value < 0.05.

### Analysis of single-cell RNA-sequencing (scRNA-seq) Data from Day 9 and 11 CVPs Derived from HS1001 and H1 hESCs Cell Lines

We generated four independent single-cell RNA-sequencing datasets with day 9 and day 11 cardiovascular progenitor cells derived from HS1001 and H1 cells respectively. H1 and HS1001 cells were cultured for 9 and 11 days with the laminin-protocol previously described to generate four independent sets of cardiac progenitor cells. In each, RNA was isolated using a 10x genomics kit and sequenced using the Illumina Hi-Seq3000 sequencing platform.

The reads were mapped to the human genome (Ensembl version 90) and quantified using Cell Ranger 2.1.1 (10x Genomics). We discarded genes with zero counts in all cells. We provided to Cell Ranger a custom built reference transcriptome generated by filtering the Ensembl transcriptome for the gene biotypes: protein coding, lincRNA and antisense. Cell Ranger was run with the expect number of cells parameter (expect-cells) set to 1000. Cell Ranger output matrices (i.e. genes.tsv and barcodes.tsv) were then input into R and genes with zero counts in all cells were discarded. For each sample (e.g. HS1001-derived CVPs at day 9) we carried out an independent cell quality control. We removed: (a) cells with very low and very high library size (i.e. cells below and above the 5^th^ and 99^th^ percentiles of the total cell library size respectively), (b) cells with low number of detected genes (i.e. cells below the 5^th^ percentile of the total number of detected genes cell distribution), (c) cells with more than 10% of their total gene count coming from mitochondrial genes. Then, all the remaining cells from the four samples were combined and we carried out five gene quality control steps: (a) we only kept “detectable” genes, defined as genes detected with more than one transcript in at least 1% of the total cells, (b) we removed genes with low average expression in the data (i.e. genes with an average expression below 0.01, this cutoff was set based on the total distribution of average gene expression across all cells and all genes), (c) we removed genes with a high dropout rate using M3Drop 3.09 R package ^73^. We used the M3Drop false discovery rate (FDR) to rank all genes from low to high FDR and then removed the bottom 25% of genes (i.e. genes with highest dropout). (d) We removed outlier genes in the gene expression distribution (i.e. “MALAT1” gene). (e) We removed genes encoded on the mitochondrial genome. After the gene quality control steps, gene counts were normalized with scran 1.8.4 R package ^74^. Scran size factors were computed from cell pools by doing a pre-clustering of the data with the quickCluster function. The output object of this function was provided to the computeSumFactors function and then we computed log-transformed normalized counts using the function *normalize* from the scater 1.8.4 R package (with default parameterizations). For each cell we computed a G2M and G1 cell cycle score by using cyclone function from the *scater* R package ^75^. We supplied to this function with the set of human cell cycle genes provided in ^76^. Then, we corrected the Log2-scran normalized genes counts for cell cycle effects by providing the counts and covariates “cell cycle G2M score” and “cell cycle G1 score” to the function removeBatchEffect from *limma* 3.36.3 R package ^77^.

Three graphical versions versions of the same t-SNE analysis were generated, one to assess the robustness of the differentiation protocol by plotting all the data combined (all the CVPs derived from different ESCs lines), Fig. 3c. In the other two t-SNE graphs we plotted CVPs from each cell line separately (Fig. 3e). The t-SNE was computed with two components and by providing the Log_2_ scran-normalized and cell cycle adjusted data from the four 10x runs to the function plotTSNE from *scater* R package ^75^. The t-SNE perplexity parameter was set to 50. Cell subpopulations were also visualized on the same t-SNE graph by mapping each cell to the expression level of known marker genes. Minimum and maximum colour was set to the minimum and maximum expression level of each of the genes plotted in the cells plotted in each graph. In addition, to test the similarity of the CVPs derived from HS1001 and H1 ECSs we computed the mean expression level (Log_2_ scran-normalized and cell cycle adjusted) for each gene in the HS1001- and H1-derived CVPs separately at day 9 and 11. Then we computed the Spearman’s ranked correlation of the CVPs from these two separate cell lines and each time point separately. We also visualized this with scatterplots, Fig. 3d.

### Human Embryonic Stem Cell Culture and Differentiation to Cardiomyocytes

All pluripotent hESCs H1 (WiCell Research Institute) and HS1001^27^ were maintained on culture plates coated overnight (4°C) with 10 μg/ml of recombinant LN-521 (Biolamina AB) in PBS. Maintenance medium Nutristem^®^ (Biological Industries) was refreshed daily ^27^. Routine monitoring of pluripotent markers (*POU5F1* and Tra1-60) by flow cytometry and genomic stability by karyotyping were performed. Cells were passaged at 80% confluence by gentle dissociation with TyrPLE (Invitrogen) at 37°C for 8 mins followed by pipetting to dissociate the single cells. The cell suspension was then collected and centrifuged at 800 rpm for 4 mins. Supernatants were discarded and the cell pellets were resuspended in 1 ml of warm Nutristem^®^ medium. Cells were split at 1:50 split ratio for H1 cells and 1:10 split ratio for HS1001 cells. Bright field images were taken with Leica microscope.

For CM differentiation, cells were seeded at 100,000 cells/cm^2^ into 24-well plates coated with the combination matrices of LN-521 and LN-221 (in ratio 1:3) at 3 μg (day 0) in PBS. This protocol was modified from Lian *et al* ^78^. At confluence (day 4), the cells were exposed to differentiation medium (RPMI 1460 with B27 supplement without insulin, Invitrogen) and 10 μM of CHIR99021 (Tocris) for 24 hours. The next day (day 5), the medium was removed and replaced with differentiation medium. At day 7, the medium was changed to differentiation medium with the addition of 5 μM of IWP2 for 2 days. Wells were replaced with differentiation medium at day 9 and day 11. Following this, medium was changed every 2 days with maintenance medium (RPMI 1460 with CTS B27 supplement with insulin (A1486701, Invitrogen) up to 94 days. Schematic presentation of the differentiation protocol is shown in Fig. 2a.

### Flow Cytometry

Specific markers were probed with antibodies using flow cytometry. Cells were dissociated and centrifuged into pellet in a microcentrifuge tube. The cells were fixed with reagent A of the Fix and Perm kit (Invitrogen) for 20 mins at room temperature, followed by a wash with 1 ml of 1% BSA in PBS and spun at 10,000 rpm for 30 sec. Subsequently, cells were permeabilized with reagent B in the Fix and Perm kit and incubated together with primary antibodies: *POU5F1* (Santa Cruz, sc-5279, 1:20), Tra1-60 (Millipore, MAB4360, 1:50), *TNNT2* (Thermo Scientific, MA-295-p1, 1:200), *MYH1* (DSHB, MF-20, 1:20) or appropriate isotype control, for 20 mins at room temperature. After incubation, the cells were washed with 1% BSA (Sigma), spun and incubated in the dark for 15 mins with the appropriate Alexa-conjugated secondary antibody (Invitrogen) at 1:1000 dilutions in 1% BSA. Finally, the cells were washed and resuspended in 300 μl of 1% BSA prior to analysis in FACS Fortessa (Becton-Dickinson). Data were analyzed using FlowJo software (Version 8). Cell gating (% expression) was done at the intersection between the isotype control and the marker expression^79^.

### Immunostaining

Cell sheets differentiated for 34 days were dissociated into single cells with TryPLE and seeded at low density onto LN-521+LN-221-coated plates. After 3 days, CMs were washed once with PBS and fixed with 4% paraformaldehyde (PFA) (Sigma) for 20 mins at 4°C. The cells were then washed with PBS and blocked with blocking buffer containing 1% BSA, 5% goat serum (Sigma) and 0.1% Triton X (Sigma) for 1 hr at room temperature. After blocking, cells were incubated with primary antibody: TNNI3 (Santa cruz, sc-15368, 1:20), NKX2-5 (Santa Cruz, sc-14033, 1:100), TNNT2 (Thermo Scientific, MS-295-p1, 1:200), *α*-actinin (Sigma, A7811, 1:500) in PBS containing 1% BSA and 5% goat serum (Gibco) at room temperature for 1 hour. Following this, antibody solution was removed and cells washed thrice with PBS on a shaker for 5 mins each. Subsequently, the cells were incubated at room temperature with Alexa-conjugated secondary antibody (Invitrogen, 1:1000) on a shaker at low speed in the dark for 1 hr. Finally, cells were washed with PBS twice (5 mins) and ProLong^®^ Gold Antifade Mountant with DAPI (Molecular probes) added to stain the nuclei. Cells were visualized immediately in Leica TCS sp8 confocal fluorescence microscope.

### Action Potential Recording

Whole cell configuration of the patch-clamp technique was used to measure the action potentials (APs) in CMs. The signal was amplified using an axon 700B patch clamp amplifier (axon instrument) and low-pass filtered at 5 kHz. Data acquisition was achieved using the Digidata 1440A (axon instrument). Data analysis and fit were performed using clamp fit 10.2 (axon instrument) and Origin 7.0 software (Origin Lab Corporation). Cells were maintained at 35°C by a temperature controller (Warner Instruments) during the recording of APs. The APs were recorded under current-clamp mode in normal Tyrode’s solution containing NaCl 140 mM, KCl 5.4 mM, CaCl_2_ 1.8 mM, MgCl_2_ 1 mM, glucose 10 mM and HEPES 10 mM, adjusted to pH 7.4 with NaOH. Pipette solution containing KCl 130 mM, NaCl 5 mM, MgCl_2_ 1 mM, MgATP 3 mM, EGTA 10 mM, and HEPES 10 mM, adjusted to pH 7.2 with KOH were utilized. AP properties include AP durations (APD) at 20%, 50% and 90% of repolarization (APD20, APD50, and APD90), AP amplitude (APA), maximal diastolic potential (MDP), and heart rates (HR) were analyzed.

The ventricular (V)- and atrial (A)-like hESC-CMs were distinguished from the nodal (N)-like hESC-CMs by higher APA and steeper dV/dt_Max_. The V-like CMs were discriminated from the A-, and N-like CMs by a longer APDs particular APD20 and a higher ratio of APD90/APD50. All CMs tested in the current study were 32~40 days post differentiation.

### Cytotoxic Drug Testing

In order to assess the viability of the CMs to cytotoxic compounds, we performed a cell viability assay using CellTiter-Glo Luminescent Cell Vaibity Assay (Promega). 96-well plates were coated with LN-521 and 15,000 CMs at day 34 were seeded into triplicate well. After 2 days of seeding, wells were washed and replenished with 90 μl of CMs maintenance medium. Cytotoxic drugs (Valinomycin (Fluka), Emetine dihydrochloride (Sigma), Doxazosin mesylate (Sigma) and Staurosporine (AG Scientific)) were prepared in 10X maintenance medium containing 10% DMSO (Sigma). Following that, 10 μl of each compound dilution was added to triplicate wells bringing the final volume to 100 μl in 1% DMSO for 24 hrs. Cellular ATP concentrations were assessed using the assay kit as per manufacturer’s instructions. Luminescence readings were taken on the Infinite 200 PRO series microplate reader (Tecan) at 1 sec integration time.

### Microelectrode Array

The extracellular field potential duration (FPD) produced by CMs was determined by the microelectrode array (MEA) assay performed with (Microelectrode array recording system (Multi Channel Systems)). Clusters of CMs were seeded on LN521-coated MEA chips containing 60 electrodes and the HR and FPD were recorded at 37ºC. FPD were recorded at baseline, exposed to 0.1 µM of isoproterenol (Sigma) or 0.01, 0.05 and 0.1 µM of E-4031 (Sigma) and after washout. FPDs were corrected by the heart rate with Bazett formula. The FPD were analyzed and corrected by heart rates with Bazett formula. The formula to calculate corrected FPD: (cFPD) (s) = FPD(s) / √ISI(s). All CMs tested in the current study were 32~40 day post differentiation.

### Luciferase-labeled Cardiovascular Progenitors

In order to track the biodistribution and viability of the progenitors, pluripotent stem cells (H1) were first transfected with luciferase plasmid and then differentiated into progenitors. Luciferase labeled CVPs were generated with 3^rd^ generation, replication incompetent RediFect Red-Fluc-Puro lentiviral particles (Perkin Elmer). 96-well plates were coated with LN-521 and 10,000 pluripotent H1 cells were seeded into each well with Nutristem^®^ medium. Next day, medium was replaced with Nutristem^®^ medium containing 4 μg/ml of hexadimethnine bromide (Sigma). Viral particles at 10 multiplicity of infection (MOI) were added directly to cells and incubated for 24 hrs. Following that, medium was replaced with fresh Nutristem^®^ for another 24 hrs. To select the successfully transfected cells, Nutristem^®^ containing 200 ng/ml of puromycin were added to eliminate cells without lentiviral particles. Finally, we assayed for the expression of luciferase with D-luciferin (Perkin Elmer) in these puromycin resistant cells. D-luciferin was diluted 1:200 in Nutristem^®^ and added into the wells. Plates were incubated at 37°C for 20 mins and luminescence signals were taken on Infinite 200 PRO series microplate reader (Tecan) at 1 sec integration time. Pluripotency of these luciferase labeled H1 cells were confirmed with flow cytomtery and cells frozen for long-term storage. The cells were differentiated according to previously described CM differentiation protocol until day 9 and day 11. The obtained progenitors were then used for in vivo experiments.

### Animals

Male CrTac:NCr-Foxn1^nu^ nude mice (8-10 weeks old, 20-25g) were purchased from InVivos, Singapore, and used for all experiments. Mice were maintained in individually ventilated cages (IVC) with filter tops and received standard diet, water and libitum. All the procedures involving animal handling were performed with prior approval and in accordance with the protocols and guidelines of Institutional Animal Care and Use Committee (IACUC), National University of Singapore.

### Surgical Procedures

Myocardial ischemia reperfusion (I/R) injury was induced as previously described ^80,81^ with modification. Briefly, mice were anaesthetized with a mixture of 0.5 mg/kg medetomidine (Pfizer Animal Health), 5.0 mg/kg Dormicum (Sciencelab) and 0.05 mg/kg Fentanyl (Pfizer Pharmaceuticals Group) and ventilated with 100% oxygen with an automated ventilator (Harvard Apparatus Inc). A core body temperature of approximately 37°C was maintained during surgery by an automatic heating blanket and continuous monitoring with a rectal thermometer. Myocardial I/R injury was induced by surgical ligation of the left coronary artery for 30 mins and releasing the ligature afterwards. Myocardial ischemia and reperfusion were confirmed by bleaching of the myocardium (and ventricular tachyarrhythmia) during ischemia and brief hyperemia in the reperfused myocardium, respectively. Five minutes before reperfusion (ligature release), 10 μL RPMI-1640 medium control or medium containing 1×10^6^ luciferase labeled CVPs (day 9 or day 11) were divided at two injection sites within the infarct zone. Mice were recovered by subcutaneous administration of 2.5 mg/kg Atipamezole (Pfizer Animal Health) and 0.5 mg/kg Flumazenil (Sagent Pharmaceuticals), followed by 0.1 mg/kg buprenorphine (Hospira Inc.) twice per day up to 3 days for analgesia. The surgeon was blinded to all cell types and medium treatments.

### Infarct Size and Area at Risk

Infarct size (area of dead myocardial tissue) and area at risk (area of ischemic myocardial tissue) were determined as previously described ^80^ with modification. In brief, 8 – 10 mice per group were anesthetized 24 hours after myocardial I/R injury and the left coronary artery was occluded again at the same level (marked by the ligature left in place after myocardial I/R injury). To determine the area at risk the heart was injected with 4% Evans Blue dye (Sigma) via the thoracic aorta. The non-ischemic tissue stained blue while the area at risk was pale. After heart extraction, the left ventricle was isolated and snap frozen in liquid nitrogen for 10 min then sliced into 5 cross sections of 1 mm thick. Heart sections were incubated at 37°C for 10 min in 1% triphenyltetrazolium chloride (Sigma) and then transferred to 4% PFA for imaging. Viable tissue is stained red while the infarcted tissue became transparent. Infarct size, area at risk and total left ventricle area were measured with ImageJ software (version 1.47c). Infarct size was expressed as percentage of area at risk and the area at risk was expressed as percentage of total left ventricle.

### Teratoma Formation Assay

To assess the *in vivo* safety of CVPs, we performed a teratoma formation assay in nude mice (8-10 weeks old, 20-25g). Single pluripotent hESCs or CVPs (5 million cells) were resuspended in 100 μl of Matrigel^TM^ (Corning) and injected intramuscularly into the hind limb muscle for 8 weeks. Samples of injected tissue were excised, fixed in formalin and routinely processed through paraffin and staining with Hematoxylin and Eosin. Images were examined under light microscopy (Olympus) by an ACVP board-certified veterinary pathologist.

### *In Vivo* and *ex Vivo* Bioluminescent Measurements

Mice received 150 mg/kg D-luciferin (Perkin Elmer) via intraperitoneal injection prior to anesthesia with 3% isoflurane (Baxter). Five minutes after injection of luciferin, the mice were placed on the heated imaging stage of the IVIS Spectrum (Perkin Elmer) in the supine position. Anesthesia was maintained with 1.5% isoflurane and 100% oxygen at 1 L/min flow during the imaging procedure. For *ex vivo* imaging to visualize the transplanted cells in situ, the extracted heart was placed on the imaging platform. Default bioluminescent settings of Living Image version 4.5 were used. Regions of interest were placed on the 2D bioluminescent image to take all bioluminescent signal and total photons emitted from the heart area were automatically calculated. Mice that had lost the luciferase signal during the 12 weeks follow-up were excluded from further analyses. A total of 4 out of 13 mice from day 9 progenitor treated group and 1 out of 8 mice from day 11 progenitor treatment were excluded.

### Myocardial Infarct Scar Analysis

To analyze fibrosis formation after myocardial I/R injury (week 12), left ventricles were fixed with 4% PFA, embedded in paraffin and sectioned in 5 μm slices. After deparaffinization and rehydration with graded concentrations of ethanol to deionized water, the sections were stained with Picrosirius Red solution (0.1 g Direct Red 80 in 100ml Picric acid solution, Sigma) for 30 mins. The sections were washed with 0.2N hydrochloric acid (Sigma) then mounted with Entellan (Merck Millipore). Staining was imaged under a Nikon Eclipse Ti inverted microscope and analyzed with NIS-Element AR Analysis software 4.5 version (Nikon Instruments Inc.). A total of 5-7 sections were analyzed per heart (n = 3) with a 100 μM distance between them. Data were analyzed by one-way ANOVA with LSD post hoc analysis.

### Assessment of Angiogenesis

Paraffin embedded heart tissue was cut into 5 μm sections for immunofluorescence staining at week 12 post-surgery. To reduce non-specific background staining, tissue sections were incubated in blocking buffer containing 10% goat serum (ThermoFisher) with for 1 hour at room temperature. Heart tissue was stained with Biotinylated Griffonia Simplicifolia Lectin I (GSL I) isolectin B4 (Vector Laboratories) in 10% goat serum (ThermoFisher) blocking buffer at 4°C overnight followed by incubation with Alexa Fluor 568 conjugated streptavidin (ThermoFisher Scientific) to visualize the microvessels, Fluorescein wheat germ agglutinin (WGA) (Vector Laboratories) was used to stain cell membranes. The sections were mounted with Fluoroshield Mounting Medium with DAPI (ab104139, Abcam). Staining was imaged under a Nikon Eclipse Ti inverted microscope (Nikon Instruments Inc.) and analyzed by NIS-Element AR Analysis software 4.5 version (Nikon Instruments Inc.). A total of 6-12 sections were analyzed per heart (n = 3) with a 100 μM distance between them. Data were analyzed by one-way ANOVA with LSD post hoc analysis.

### Immunohistochemistry

Mouse hearts were coronally dissected at the middle of the infracted area using a clean scalpel blade per cut at week 12 post MI. Heart tissues were fixed in 4% PFA overnight and followed with 30% sucrose in PBS at 4°C. Tissues were embedded in optimal cutting temperature compound and sectioned. Tissue sections were washed with 1 x PBS thrice for 5 mins each and antigen retrieval with proteinase K (DAKO) for 5 mins. After which, sections were washed thrice with 1 X PBS for 5 mins each and incubated with 0.2% Triton X-100 (Sigma) for 10 mins. Sections were washed and blocked with 5% donkey serum (Sigma) for 30 mins. Sections were then incubated with specific primary antibody: anti-TNNT2 (Thermo scientific, MS-295-p1, 1:100), anti-Ku80 (Abcam, ab80592, 1:100), anti-CX43 (BD, 610061, 1:100), 4°C overnight.

Next day, for immunofluoresecence staining, sections were washed thrice with 1 X PBS for 7 mins each. Secondary antibody was added for 1 hr and washed thrice with 1 X PBS for 7 mins each. Finally, slides were mounted with ProLong Gold antifade mountant with DAPI (P36931, Life Technologies). Slides were examined using Leica confocal microscope.

For 3,3′-Diaminobenzidine staining, sections were incubated with peroxidase blocking solution (DAKO) for 10 mins and wash with PBS. Secondary antibody was added for 1 hr followed by washing. After which, sections were incubated with DAB solution (DAKO) for 1-3 mins and counter stained with hematoxylin for 2 mins. Finally, all sections were dehydrated with ethanol and Clearene (Leica). The slides were mounted with Eukitt quick-hardening mounting media (Fluka) and examined using light microscope (Olympus).

### Cardiac Function Assessment

Cardiac function was assessed as previously described^81^. In brief, echocardiogram was performed with a high frequency ultrasound system Vevo^®^ 2100 (Visualsonics) on mice under general anesthesia (1-1.5% isoflurane, Baxter) at baseline, week 1, 4, 8 and 12 after myocardial I/R injury. Body temperature was monitored with a rectal probe and maintained at 36-37°C. Volumes and functional parameters were measured from parasternal long-axis B-mode images (LV trace mode). Long-axis B-mode images were also used for speckle-tracking strain analysis (both longitudinal and radial strain analysis) with the VevoStrain software (Visualsonics) as previously described ^82^. All the data were analyzed by an experienced researcher in a blinded fashion with Vevo^®^ 2100 software, version 1.7.0 (Visualsonics). Representative results at week 8 post MI were shown in Fig. 5.

### Statistical Analysis

Comparisons between groups in time were performed using two-way ANOVA with Bonferroni *post hoc* analysis unless otherwise stated. Values were reported as Mean ± SEM. P-value ≤ 0.05 was considered statistically significant. Data were analyzed with SPSS software (IBM^®^ SPSS^®^ Statistics version 22.0) and GraphPad Prism (Version 7.0a).

## ACKNOWLEDGMENTS

We thank Professor David Silver and Assistant Professor Manvendra Singh for reading the manuscript. We also thank the Duke NUS Genome Biology Facility and Advanced Bioimaging Core, Singapore General Health Services for their RNA sequencing and for confocal microscopy services respectively.

## AUTHOR CONTRIBUTIONS

L.Y. designed and performed experiments, analyzed data and wrote the manuscript. J.W.W designed, performed the animal experiments and wrote the manuscript. J.W.W., X.W. and S.Y.C. performed the *in vivo* experiments and histology staining, analyzed and interpreted the data. A.M.M. conceptualized and analyzed all the RNA sequencing and wrote the manuscript. Y.S. cloned and generated the LN-221 protein. L.Y.C. performed all the heart histology, immunofluorescence staining and microscopic data analysis. N.H., M.O. and S.G. assisted the conceptualization and analyses of RNA sequencing, H.W. performed the electrophysiology measurements and R.B. assisted in histology analysis. O.H. generated the HS1001 cell line and E.P. worked with A.M.M. on the RNA sequencing analyses. S.C. and D.P.V.d.K. contributed to the heart experiments, data analyses and helped with the editing of the manuscript. K.T. conceived and supervised the overall project and wrote the manuscript. All authors vetted and approved the final manuscript.

## REFERENCES

1 Bergmann, O. et al. Evidence for cardiomyocyte renewal in humans. Science (New York, N.Y.) 324, 98–102, doi:10.1126/science.1164680 (2009).

2 Tzahor, E. & Poss, K. D. Cardiac regeneration strategies: Staying young at heart. Science (New York, N.Y.) 356, 1035–1039, doi:10.1126/science.aam5894 (2017).

3 Quyyumi, A. A. et al. PreSERVE-AMI: A Randomized, Double-Blind, Placebo-Controlled Clinical Trial of Intracoronary Administration of Autologous CD34+ Cells in Patients With Left Ventricular Dysfunction Post STEMI. Circulation research 120, 324–331, doi:10.1161/CIRCRESAHA.115.308165 (2017).

4 Cahill, T. J., Choudhury, R. P. & Riley, P. R. Heart regeneration and repair after myocardial infarction: translational opportunities for novel therapeutics. Nat Rev Drug Discov 16, 699–717, doi:10.1038/nrd.2017.106 (2017).

5 Zhang, J. et al. Functional cardiomyocytes derived from human induced pluripotent stem cells. Circ Res 104, e30–41, doi:10.1161/CIRCRESAHA.108.192237 (2009).

6 Fernandes, S. et al. Comparison of Human Embryonic Stem Cell-Derived Cardiomyocytes, Cardiovascular Progenitors, and Bone Marrow Mononuclear Cells for Cardiac Repair. Stem Cell Reports 5, 753–762, doi:10.1016/j.stemcr.2015.09.011 (2015).

7 Shiba, Y. et al. Human ES-cell-derived cardiomyocytes electrically couple and suppress arrhythmias in injured hearts. Nature 489, 322–325, doi:10.1038/nature11317 (2012).

8 Chong, J. J. et al. Human embryonic-stem-cell-derived cardiomyocytes regenerate non-human primate hearts. Nature 510, 273–277, doi:10.1038/nature13233 (2014).

9 Weinberger, F. et al. Cardiac repair in guinea pigs with human engineered heart tissue from induced pluripotent stem cells. Science translational medicine 8, 363ra148, doi:10.1126/scitranslmed.aaf8781 (2016).

10 Liu, Y. W. et al. Human embryonic stem cell-derived cardiomyocytes restore function in infarcted hearts of non-human primates. Nat Biotechnol 36, 597–605, doi:10.1038/nbt.4162 (2018).

11 Nam, Y. J. et al. Reprogramming of human fibroblasts toward a cardiac fate. Proc Natl Acad Sci U S A 110, 5588–5593, doi:10.1073/pnas.1301019110 (2013).

12 Qian, L. et al. In vivo reprogramming of murine cardiac fibroblasts into induced cardiomyocytes. Nature 485, 593–598, doi:10.1038/nature11044 (2012).

13 Warnes, G. R. et al. gplots: Various R Programming Tools for Plotting Data (2016).

14 Schussler-Lenz, M. et al. Cell-based therapies for cardiac repair: a meeting report on scientific observations and European regulatory viewpoints. Eur J Heart Fail 18, 133–141, doi:10.1002/ejhf.422 (2016).

15 Murry, C. E., Chong, J. J. & Laflamme, M. A. Letter by murry et Al regarding article, “embryonic stem cell-derived cardiac myocytes are not ready for human trials”. Circulation research 115, e28–29, doi:10.1161/circresaha.114.305042 (2014).

16 Sussman, M. A. & Puceat, M. Response to letter regarding article, “embryonic stem cell-derived cardiac myocytes are not ready for human trials”. Circulation research 115, e30–31, doi:10.1161/circresaha.114.305341 (2014).

17 Orkin, R. W. et al. A murine tumor producing a matrix of basement membrane. The Journal of experimental medicine 145, 204–220 (1977).

18 Kleinman, H. K. et al. Isolation and characterization of type IV procollagen, laminin, and heparan sulfate proteoglycan from the EHS sarcoma. Biochemistry 21, 6188–6193 (1982).

19 Lu, T. Y. et al. Repopulation of decellularized mouse heart with human induced pluripotent stem cell-derived cardiovascular progenitor cells. Nature communications 4, 2307, doi:10.1038/ncomms3307 (2013).

20 Melkoumian, Z. et al. Synthetic peptide-acrylate surfaces for long-term self-renewal and cardiomyocyte differentiation of human embryonic stem cells. Nat Biotechnol 28, 606–610, doi:10.1038/nbt.1629 (2010).

21 Nagaoka, M., Si-Tayeb, K., Akaike, T. & Duncan, S. A. Culture of human pluripotent stem cells using completely defined conditions on a recombinant E-cadherin substratum. BMC developmental biology 10, 60, doi:10.1186/1471-213X-10-60 (2010).

22 Braam, S. R. et al. Recombinant vitronectin is a functionally defined substrate that supports human embryonic stem cell self-renewal via alphavbeta5 integrin. Stem cells 26, 2257–2265, doi:10.1634/stemcells.2008-0291 (2008).

23 Domogatskaya, A., Rodin, S. & Tryggvason, K. Functional diversity of laminins. Annual review of cell and developmental biology 28, 523–553, doi:10.1146/annurev-cellbio-101011-155750 (2012).

24 Miner, J. H. & Yurchenco, P. D. Laminin functions in tissue morphogenesis. Annual review of cell and developmental biology 20, 255–284, doi:10.1146/annurev.cellbio.20.010403.094555 (2004).

25 Domogatskaya, A., Rodin, S., Boutaud, A. & Tryggvason, K. Laminin-511 but not −332, −111, or −411 enables mouse embryonic stem cell self-renewal in vitro. Stem cells 26, 2800–2809, doi:10.1634/stemcells.2007-0389 (2008).

26 Rodin, S. et al. Long-term self-renewal of human pluripotent stem cells on human recombinant laminin-511. Nature Biotechnology 28, 611–U102, doi:10.1038/nbt.1620 (2010).

27 Rodin, S. et al. Clonal culturing of human embryonic stem cells on laminin-521/E-cadherin matrix in defined and xeno-free environment. Nature communications 5, 3195, doi:10.1038/ncomms4195 (2014).

28 Nguyen, M. T. et al. Differentiation of Human Embryonic Stem Cells to Endothelial Progenitor Cells on Laminins in Defined and Xeno-free Systems. Stem Cell Reports 7, 802–816, doi:10.1016/j.stemcr.2016.08.017 (2016).

29 Sigmundsson, K. et al. Culturing functional pancreatic islets on alpha5-laminins and curative transplantation to diabetic mice. Matrix biology : journal of the International Society for Matrix Biology 70, 5–19, doi:10.1016/j.matbio.2018.03.018 (2018).

30 Guyette, J. P. et al. Bioengineering Human Myocardium on Native Extracellular Matrix. Circulation research 118, 56–72, doi:10.1161/CIRCRESAHA.115.306874 (2016).

31 Heinig, M. et al. Natural genetic variation of the cardiac transcriptome in non-diseased donors and patients with dilated cardiomyopathy. Genome biology 18, 170, doi:10.1186/s13059-017-1286-z (2017).

32 Consortium, G. T. Human genomics. The Genotype-Tissue Expression (GTEx) pilot analysis: multitissue gene regulation in humans. Science (New York, N.Y.) 348, 648–660, doi:10.1126/science.1262110 (2015).

33 Kortesmaa, J., Yurchenco, P. & Tryggvason, K. Recombinant laminin-8 (alpha(4)beta(1)gamma(1)). Production, purification,and interactions with integrins. The Journal of biological chemistry 275, 14853–14859 (2000).

34 Doi, M. et al. Recombinant human laminin-10 (alpha5beta1gamma1). Production, purification, and migration-promoting activity on vascular endothelial cells. The Journal of biological chemistry 277, 12741–12748, doi:10.1074/jbc.M111228200 (2002).

35 Oram, S. H. et al. Bivalent promoter marks and a latent enhancer may prime the leukaemia oncogene LMO1 for ectopic expression in T-cell leukaemia. Leukemia 27, 1348–1357, doi:10.1038/leu.2013.2 (2013).

36 Fritsch, A. mcclust: Process an MCMC Sample of Clusterings (2012).

37 Bolstad, B. M. preprocessCore: A collection of pre-processing functions (2016).

38 Azhar, M. et al. Transforming growth factor Beta2 is required for valve remodeling during heart development. Developmental dynamics : an official publication of the American Association of Anatomists 240, 2127–2141, doi:10.1002/dvdy.22702 (2011).

39 Marvin, M. J., Di Rocco, G., Gardiner, A., Bush, S. M. & Lassar, A. B. Inhibition of Wnt activity induces heart formation from posterior mesoderm. Genes & development 15, 316–327, doi:10.1101/gad.855501 (2001).

40 Esham, M., Bryan, K., Milnes, J., Holmes, W. B. & Moncman, C. L. Expression of nebulette during early cardiac development. Cell Motil Cytoskeleton 64, 258–273, doi:10.1002/cm.20180 (2007).

41 Durinck, S., Spellman, P. T., Birney, E. & Huber, W. Mapping identifiers for the integration of genomic datasets with the R/Bioconductor package biomaRt. Nature protocols 4, 1184–1191, doi:10.1038/nprot.2009.97 (2009).

42 Loh, Y. H. et al. The Oct4 and Nanog transcription network regulates pluripotency in mouse embryonic stem cells. Nature genetics 38, 431–440, doi:10.1038/ng1760 (2006).

43 Zheng, Y. et al. Metallothionein 1H (MT1H) functions as a tumor suppressor in hepatocellular carcinoma through regulating Wnt/beta-catenin signaling pathway. BMC cancer 17, 161, doi:10.1186/s12885-017-3139-2 (2017).

44 Boles, N. C. et al. NPTX1 regulates neural lineage specification from human pluripotent stem cells. Cell Rep 6, 724–736, doi:10.1016/j.celrep.2014.01.026 (2014).

45 Mummery, C. L. et al. Differentiation of human embryonic stem cells and induced pluripotent stem cells to cardiomyocytes: a methods overview. Circulation research 111, 344–358, doi:10.1161/circresaha.110.227512 (2012).

46 Moretti, A. et al. Patient-specific induced pluripotent stem-cell models for long-QT syndrome. The New England journal of medicine 363, 1397–1409, doi:10.1056/NEJMoa0908679 (2010).

47 Ma, J. et al. High purity human-induced pluripotent stem cell-derived cardiomyocytes: electrophysiological properties of action potentials and ionic currents. American journal of physiology. Heart and circulatory physiology 301, H2006–2017, doi:10.1152/ajpheart.00694.2011 (2011).

48 He, J. Q., Ma, Y., Lee, Y., Thomson, J. A. & Kamp, T. J. Human embryonic stem cells develop into multiple types of cardiac myocytes: action potential characterization. Circulation research 93, 32–39, doi:10.1161/01.RES.0000080317.92718.99 (2003).

49 Zwi, L. et al. Cardiomyocyte differentiation of human induced pluripotent stem cells. Circulation 120, 1513–1523, doi:10.1161/CIRCULATIONAHA.109.868885 (2009).

50 Hoekstra, M., Mummery, C. L., Wilde, A. A., Bezzina, C. R. & Verkerk, A. O. Induced pluripotent stem cell derived cardiomyocytes as models for cardiac arrhythmias. Front Physiol 3, 346, doi:10.3389/fphys.2012.00346 (2012).

51 Moretti, A. et al. Multipotent embryonic isl1+ progenitor cells lead to cardiac, smooth muscle, and endothelial cell diversification. Cell 127, 1151–1165, doi:10.1016/j.cell.2006.10.029 (2006).

52 Wu, S. M. Mesp1 at the heart of mesoderm lineage specification. Cell Stem Cell 3, 1–2, doi:10.1016/j.stem.2008.06.017 (2008).

53 Cai, C. L. et al. Isl1 identifies a cardiac progenitor population that proliferates prior to differentiation and contributes a majority of cells to the heart. Dev Cell 5, 877–889 (2003).

54 Hensman, J., Rattray, M. & Lawrence, N. D. Fast Nonparametric Clustering of Structured Time-Series. IEEE transactions on pattern analysis and machine intelligence 37, 383–393, doi:10.1109/TPAMI.2014.2318711 (2015).

55 Lian, X. et al. Robust cardiomyocyte differentiation from human pluripotent stem cells via temporal modulation of canonical Wnt signaling. Proceedings of the National Academy of Sciences of the United States of America 109, E1848–1857, doi:10.1073/pnas.1200250109 (2012).

56 Burridge, P. W. et al. Chemically defined generation of human cardiomyocytes. Nature methods 11, 855–860, doi:10.1038/nmeth.2999 (2014).

57 Kadota, S., Pabon, L., Reinecke, H. & Murry, C. E. In Vivo Maturation of Human Induced Pluripotent Stem Cell-Derived Cardiomyocytes in Neonatal and Adult Rat Hearts. Stem Cell Reports 8, 278–289, doi:10.1016/j.stemcr.2016.10.009 (2017).

58 Takuya Kuroda, S. Y., Satoko Matsuyama, Keiko Tano, Shinji Kusakawa, Yoshiki Sawa, Shin Kawamata, Yoji Sato. Highly sensitive droplet digital PCR method for detection of residual undifferentiated cells in cardiomyocytes derived from human pluripotent stem cells Regenerative Therapy 2, 17–23 (2015).

59 Minami, I. et al. A small molecule that promotes cardiac differentiation of human pluripotent stem cells under defined, cytokine- and xeno-free conditions. Cell Rep 2, 1448–1460, doi:10.1016/j.celrep.2012.09.015 (2012).

60 Bassat, E. et al. The extracellular matrix protein Agrin promotes heart regeneration in mice. Nature, doi:10.1038/nature22978 (2017).

61 Zhu, K. et al. Lack of Remuscularization Following Transplantation of Human Embryonic Stem Cell-Derived Cardiovascular Progenitor Cells in Infarcted Nonhuman Primates. Circulation research 122, 958–969, doi:10.1161/circresaha.117.311578 (2018).

62 Lawrence, M. et al. Software for computing and annotating genomic ranges. PLoS computational biology 9, e1003118, doi:10.1371/journal.pcbi.1003118 (2013).

63 Love, M. I., Huber, W. & Anders, S. Moderated estimation of fold change and dispersion for RNA-seq data with DESeq2. Genome biology 15, 550, doi:10.1186/s13059-014-0550-8 (2014).

64 Reimand, J., Kolde, R. & Arak, T. gProfileR: Interface to the ’g:Profiler’ Toolkit (2016).

65 Yu, G. et al. GOSemSim: an R package for measuring semantic similarity among GO terms and gene products. Bioinformatics 26, 976–978, doi:10.1093/bioinformatics/btq064 (2010).

66 Edgar, R., Domrachev, M. & Lash, A. E. Gene Expression Omnibus: NCBI gene expression and hybridization array data repository. Nucleic acids research 30, 207–210 (2002).

67 Martin, M. Cutadapt removes adapter sequences from high-throughput sequencing reads. EMBnet.journal 17, 10–12, doi:http://dx.doi.org/10.14806/ej.17.1.200 (2011).

68 Dobin, A. et al. STAR: ultrafast universal RNA-seq aligner. Bioinformatics 29, 15–21, doi:10.1093/bioinformatics/bts635 (2013).

69 Li, B. & Dewey, C. N. RSEM: accurate transcript quantification from RNA-Seq data with or without a reference genome. BMC bioinformatics 12, 323, doi:10.1186/1471-2105-12-323 (2011).

70 Team, R. C. R: A Language and Environment for Statistical Computing (Vienna, Austria, 2016).

71 Takahashi, K. et al. Induction of pluripotent stem cells from adult human fibroblasts by defined factors. Cell 131, 861–872, doi:10.1016/j.cell.2007.11.019 (2007).

72 Yang, X., Pabon, L. & Murry, C. E. Engineering adolescence: maturation of human pluripotent stem cell-derived cardiomyocytes. Circulation research 114, 511–523, doi:10.1161/CIRCRESAHA.114.300558 (2014).

73 Andrews, T. M3Drop: Michaelis-Menten Modelling of Dropouts in single-cell RNASeq. R package version 3.09.00 (2017).

74 Lun, A. T., McCarthy, D. J. & Marioni, J. C. A step-by-step workflow for low-level analysis of single-cell RNA-seq data with Bioconductor. F1000Res 5, 2122, doi:10.12688/f1000research.9501.2 (2016).

75 McCarthy, D. J., Campbell, K. R., Lun, A. T. & Wills, Q. F. Scater: pre-processing, quality control, normalization and visualization of single-cell RNA-seq data in R. Bioinformatics 33, 1179–1186, doi:10.1093/bioinformatics/btw777 (2017).

76 Scialdone, A. et al. Computational assignment of cell-cycle stage from single-cell transcriptome data. Methods 85, 54–61, doi:10.1016/j.ymeth.2015.06.021 (2015).

77 Ritchie, M. E. et al. limma powers differential expression analyses for RNA-sequencing and microarray studies. Nucleic acids research 43, e47, doi:10.1093/nar/gkv007 (2015).

78 Lian, X. et al. Directed cardiomyocyte differentiation from human pluripotent stem cells by modulating Wnt/beta-catenin signaling under fully defined conditions. Nature protocols 8, 162–175, doi:10.1038/nprot.2012.150 (2013).

79 Yap, L. Y. et al. Defining a threshold surface density of vitronectin for the stable expansion of human embryonic stem cells. Tissue engineering. Part C, Methods 17, 193–207, doi:10.1089/ten.TEC.2010.0328 (2011).

80 Arslan, F. et al. Myocardial ischemia/reperfusion injury is mediated by leukocytic toll-like receptor-2 and reduced by systemic administration of a novel anti-toll-like receptor-2 antibody. Circulation 121, 80–90, doi:10.1161/CIRCULATIONAHA.109.880187 (2010).

81 Allijn, I. E. et al. Liposome encapsulated berberine treatment attenuates cardiac dysfunction after myocardial infarction. Journal of controlled release : official journal of the Controlled Release Society 247, 127–133, doi:10.1016/j.jconrel.2016.12.042 (2017).

82 Bhan, A. et al. High-frequency speckle tracking echocardiography in the assessment of left ventricular function and remodeling after murine myocardial infarction. American journal of physiology. Heart and circulatory physiology 306, H1371–1383, doi:10.1152/ajpheart.00553.2013 (2014).

