## Supplementary for "*In vivo* generation of post-infarct mouse cardiac muscle by cardiomyocyte progenitors produced with a reproducible laminin-promoted human stem cell differentiation system"

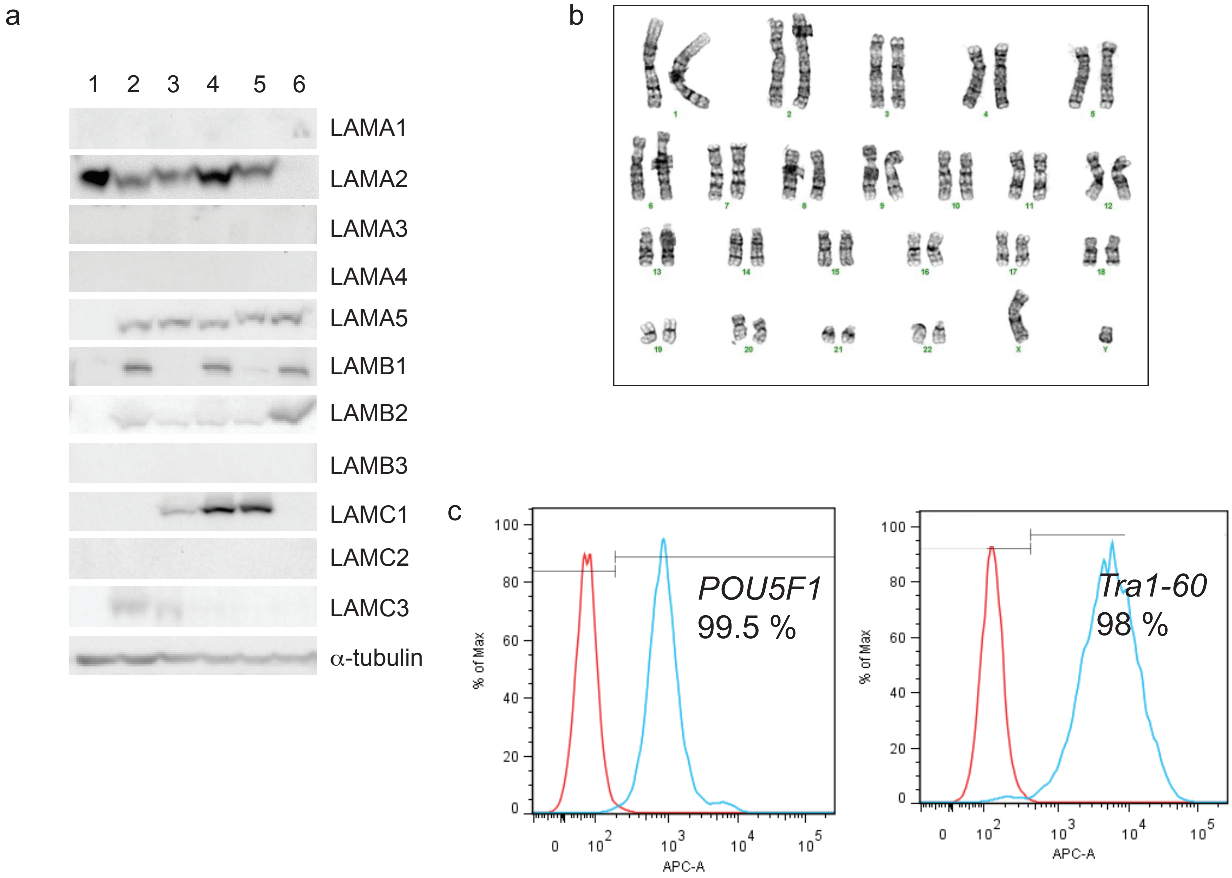


Figure S1. Expression of laminin chains in human samples and characterization of hESCs. **a** Western blot of human samples probed for individual laminin chains. **b** Chromosomal stability of H1 cells cultured on LN-521. **C**, H1 CVPs at day 9 of differentiation stained with *ISL1* antibody (Red) and DAPI (Blue). **c** Representative traces of *POU5F1* and Tra1-60 flow cytometry in HS1001 cells. Measurement revealed a high (99.5% and 98% respectively) population of pluripotent stem cells. Red trace = Isotype control and blue trace = *POU5F1* or Tra1-60 positive.

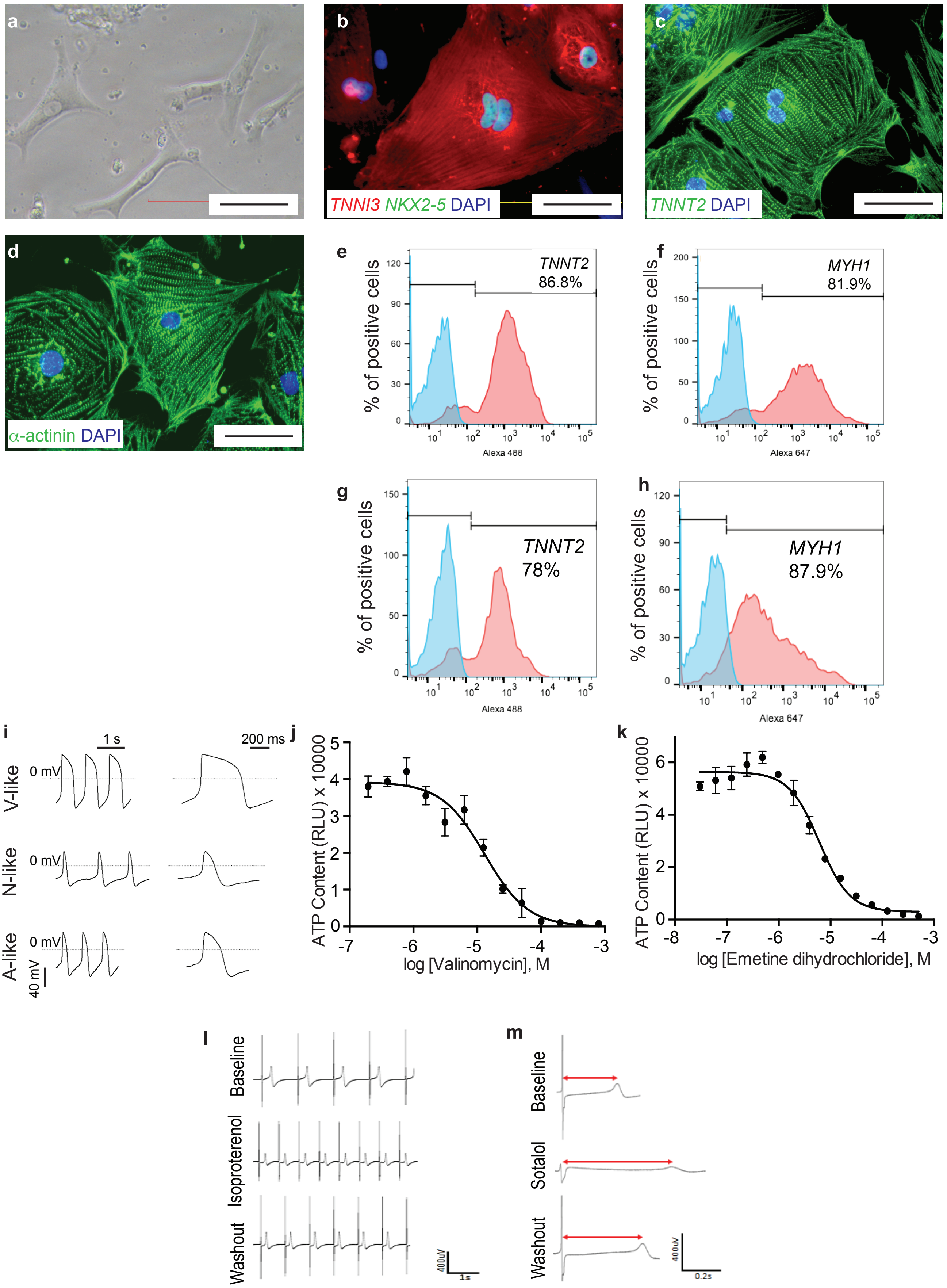

a
b
TNNI3 NKX2-5 DAPI
c
TNNT2 DAPI
d
α-actinin DAPI
e
TNNT2
86.8%
% of positive cells
Alexa 488
f
MYH1
81.9%
% of positive cells
Alexa 647
g
TNNT2
78%
% of positive cells
Alexa 488
h
MYH1
87.9%
% of positive cells
Alexa 647
i
1 s
200 ms
V-like
N-like
A-like
0 mV
40 mV
j
ATP Content (RLU) x 10000
log [Valinomycin], M
k
ATP Content (RLU) x 10000
log [Emetine dihydrochloride], M
l
Baseline
Isoproterenol
Washout
400uV
1s
m
Baseline
Sotalol
Washout
400uV
0.2s

Figure S2. Characterization hESC-derived Cardiomyocytes after 34-Day Differentiation *in Vitro*. **a** Representative bright field photomicrograph of dissociated CMs exhibiting typical spindle-shaped morphology of the cells. **b** Overlap immunofluorescence staining of cardiac specific Troponin I (*TNNI3*) in red and transcription factor *NKX2-5* in green. Nuclei were stained with DAPI (blue). **c** Overlap immunofluorescence staining of cardiac specific Troponin T (*TNNT2*) in green and nuclei stained with DAPI (blue). **d** Overlap immunofluorescence staining of α-actinin antibody in green reveals cytoplasmic sacromeric striations and nucleus stained with DAPI (blue). The images were taken at 20X magnifications. Scale bar = 100 μM. **e** Representative traces of *TNNT2* (86.8%) and isotype control (blue) as measured by flow cytometry in H1 cells. **f** Representative traces of *MYH1* (81.9%) and isotype control (blue) by flow cytometry in H1 cells. **g** Representative traces of *TNNT2* (78 %) and **h** *MYH1* (87.9 %) flow cytometry measurement by flow cytometry in HS1001 derived CMs Red trace = Isotype control and blue trace = *TNNT2* or *MYH1* positive. **i** Representative traces of action potentials that represent Ventricular (V)-, Nodal (N)- and Atrial (A)-like CMs. **j** Pharmacological drug testing on the CMs with valinomycin ($EC_{50}$ values of 13.1 μM). n = 3. **k** Emetine dihydrochloride ($EC_{50}$ values of 6.079 μM) as measured by cellular ATP concentration. n = 3. Action potentials and field potentials recorded in H1 CMs. The field potential duration (FPD) traces recorded at baseline, exposed to **l** 0.1 µM of Isoproterenol or **m** 50 µM of Sotalol and after washout. The red lines indicated FPD in each trace.

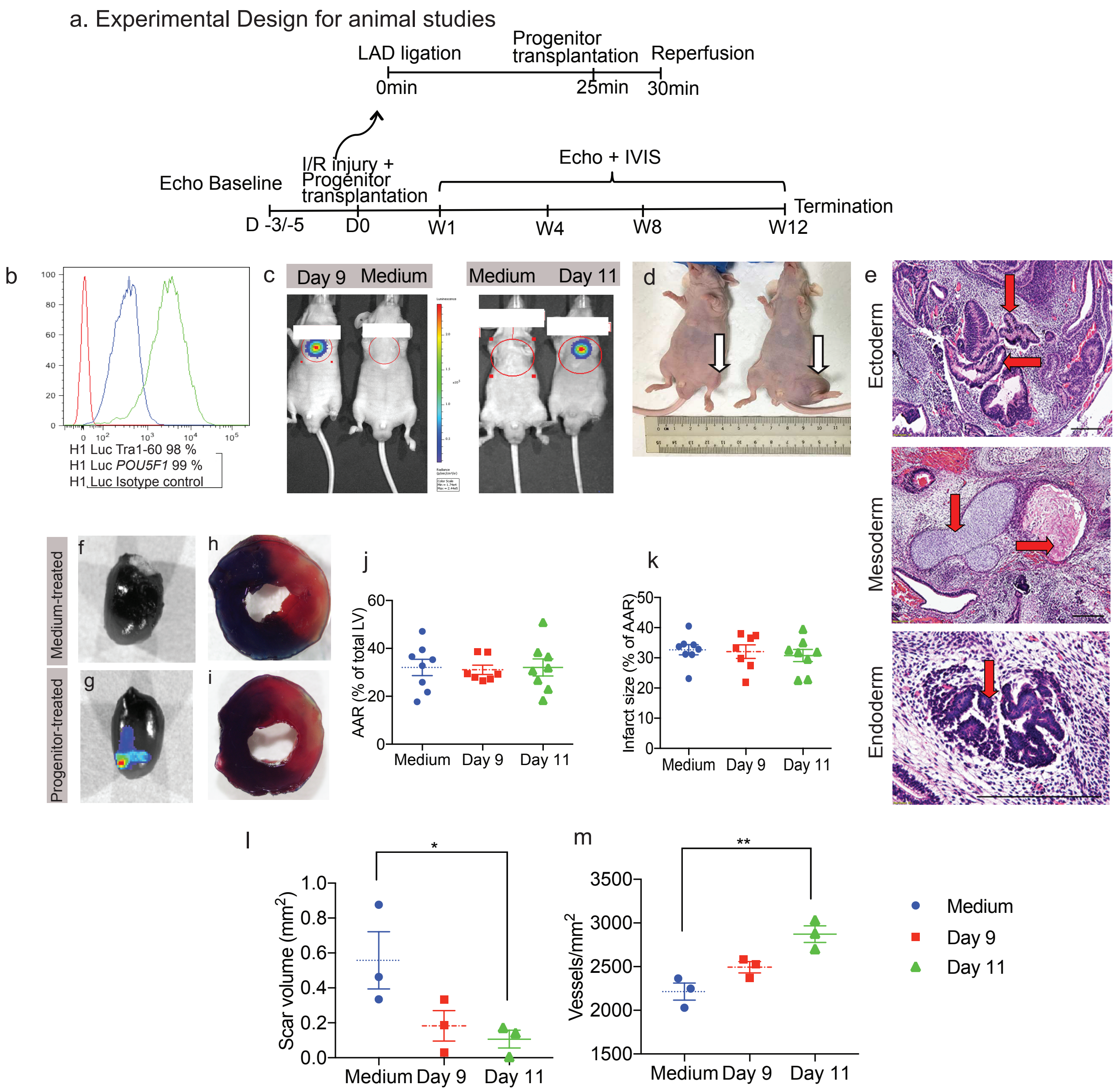
a. Experimental Design for animal studies
LAD ligation
Progenitor transplantation
Reperfusion
0min
25min
30min
Echo Baseline
I/R injury + Progenitor transplantation
Echo + IVIS
Termination
D -3/-5
D0
W1
W4
W8
W12
b
H1 Luc Tra1-60 98 %
H1 Luc POU5F1 99 %
H1 Luc Isotype control
c
Day 9
Medium
Medium
Day 11
d
e
Ectoderm
Mesoderm
Endoderm
Medium-treated
Progenitor-treated
f
g
h
i
j
AAR (% of total LV)
Medium
Day 9
Day 11
k
Infarct size (% of AAR)
Medium
Day 9
Day 11
l
Scar volume (mm²)
Medium
Day 9
Day 11
m
Vessels/mm²
Medium
Day 9
Day 11
Medium
Day 9
Day 11

Figure S3. In vivo experiments. **a** Experimental design for animal studies. **b** Flow cytometry was performed on the luciferase labeled pluripotent H1 cells. These cells showed high expression of Tra1-60 (98%) and *POU5F1* (99%). **c** A representative IVIS image from the mice at week 12 after cell transplantation showed strong biolumineiscence signal indicating long-term progenitors survival *in vivo*. Luciferase signal was only detected in the heart and not in other organs. **d** Photomicrograph of mouse injected with progenitor (left) and H1 pluripotent stem cells (right) after 8 weeks in the hindlimb muscle. **e** Hematoxylin and Eosin staining of teratoma isolated from H1 stem cells administration showing ectoderm (neural rosette), mesoderm (bone and cartilage) and endoderm (pancreas) germ layers. Scale bar = 200 mM. **f** Epiluminescence were measured using IVIS system 24 hours after myocardial I/R injury in medium-treated hearts. **g** Epiluminescence were measured using IVIS system 24 hours after myocardial I/R injury in progenitor-treated hearts. **h** Left ventricle sections were stained with TTC to quantitate the ischemic region (area at risk, AAR) in medium-treated and **i** Left ventricle sections were stained with TTC to quantitate the ischemic region (area at risk, AAR) in progenitor-treated hearts. Non-ischemic region was stained blue while ischemic region was unstained. Viable tissue was stained red and infarcted tissue was pale. **j** For each treatment group, AAR and **k** infarct size were quantified. n =7-8 per treatment group. **l** Quantification of scar volume in the infarcted area. Data were analyzed by one-way ANOVA with LSD post hoc analysis. **m** Quantification of angiogenesis in the infarcted area. Data were analyzed by one-way ANOVA with LSD post hoc analysis. Bars represent Mean ± SEM. * $p < 0.05$, ** $p < 0.01$.

Table S1 (excel format). LN-221 transcriptomic signature. Results included: 1. List of genes differentially expressed in H1 cells cultured on LN-221+LN-521 for four days when comparing against H1 cells cultured on LN-521 for four days (Benjamini-Hochberg (BH) adjusted p-value < 0.05). Log2FC, shrunken Log2-fold changes computed by DESeq2 package. BHadjP, BH adjusted p-value. Tabs 2 and 3: Functional enrichments of up and downregulated genes included in LN-221 transcriptional signature. The results include Gene Ontology GO (BP, MF and CC), KEGG and Reactome.

Table S2. Action potential parameters

| | Ventricular | Atrial | Nodal |
|---|---|---|---|
| APA, mV | 104.60±0.90 | 90.04±2.61 | 80.98±1.97 |
| Overshoot, mV | 44.12±0.81 | 40.58±2.26 | 29.81±1.12 |
| MDP, mV | -60.5±0.64 | -53.46±1.55 | -51.17±1.65 |
| APD90, ms | 446.93±17.68 | 267.60±19.42 | 187.63±15.63 |
| APD50, ms | 388.67±16.07 | 213.12±16.51 | 129.31±13.69 |
| APD20, ms | 268.56±12.97 | 142.23±13.48 | 81.76±9.93 |
| APD90/APD50 | 1.15±0.01 | 1.26±0.02 | 1.49±0.05 |
| $dV/dt_{max}$, V/s | 16.78±2.59 | 12.02±3.28 | 11.02±4.19 |
| Rate, bpm | 60.58±3.08 | 65.45±5.84 | 73.26±7.32 |
| Percentage | 64.20% | 22.22% | 13.58% |
| n | 52 | 18 | 11 |

APA, action potential amplitude; MDP, maximum diastolic potential; APD90, APD50, APD20, AP duration measured at 90%, 50% or 20% of repolarization; $dV/dt_{max}$, maximum rate of depolarization or maximal upstroke velocity; n, the cell number. Data are presented as mean ± SEM.
